# Florfenicol administration in piglets co-selects for multiple antimicrobial resistance genes

**DOI:** 10.1101/2024.06.03.597168

**Authors:** Devin B. Holman, Katherine E. Gzyl, Arun Kommadath

## Abstract

Florfenicol is a broad-spectrum phenicol antibiotic used in swine for various indications. However, information regarding its effect on the pig gut microbiome and resistome is lacking. Therefore, this study investigated those effects by treating piglets with an intramuscular injection of florfenicol at 1 and 7 days of age. Fecal samples were collected from treated (n =30) and untreated (n = 30) pigs at nine different time points up until 140 days of age and their microbiomes were profiled using both 16S rRNA gene and shotgun metagenomic sequencing. The gut microbiomes of the two groups of piglets were most dissimilar in the immediate period following florfenicol administration. These differences were driven in part by an enrichment in *Clostridium scindens*, *Enterococcus faecalis*, and *Escherichia* spp. in the florfenicol-treated piglets and *Fusobacterium* spp., *Pauljensenia hyovaginalis,* and *Ruminococcus gnavus* in the control piglets. In addition to florfenicol resistance genes including *floR*, *fexA*, and *fexB*, florfenicol also selected for genes conferring resistance to the aminoglycosides, beta-lactams, peptides, or sulfonamides up until weaning at 21 days of age. Florfenicol-resistant *Escherichia coli* isolated from these piglets were found to carry a plasmid with a *floR*, along *tet*(A), *aph(6)-Id*, *aph(3’’)-Ib*, *sul2*, and *bla*_TEM-1_/ *bla*_CMY-2_. A plasmid carrying *fexB* and *poxtA* was identified in florfenicol-resistant *Enterococcus avium*, *Enterococcus faecium*, and *E*. *faecalis* isolates from the treated piglets. This study highlights the potential for co-selection and perturbation of the gut microbial community in pre-weaned piglets administered florfenicol.

**Importance:** Antimicrobial use and resistance remain a serious challenge in food-animal production systems. Understanding how specific antimicrobials affect the gut microbiome and resistome is an important step in reducing antimicrobial use and resistance. Florfenicol is an antimicrobial used in swine production, yet very little is known about its effect on the pig gut microbiome and resistome. In this study, we administered florfenicol to piglets at 1 and 7 days of age and characterized their fecal metagenomes through to 140 days of age. Florfenicol altered the fecal microbiome and selected for many unrelated antimicrobial resistance genes up until weaning at 21 days of age. Part of this co-selection process appeared to involve an *Escherichia coli* plasmid carrying a florfenicol resistance gene along with genes conferring resistance to at least four other antimicrobial classes. These results demonstrate the potential for certain antimicrobials to co-select for multiple, unrelated antimicrobial resistance genes in pigs.

## Introduction

Antimicrobial resistance is a serious threat to both animal and human health. In commercial swine production, antimicrobials are frequently used to both treat and prevent infectious disease. However, antimicrobial use can be associated with the development and maintenance of antimicrobial resistance in pigs and may also affect the gut microbiome. In swine operations, antimicrobials are usually administered via feed, water, or intramuscular injection. Florfenicol is a broad-spectrum phenicol antibiotic that inhibits bacterial protein synthesis by binding to the 50S ribosomal subunit (1). Although florfenicol is only used in food-producing animals, it is structurally similar to chloramphenicol which is used in human medicine. Unlike some older antimicrobials such as chlortetracycline and tylosin, florfenicol is a relatively recent addition to veterinary medicine and was only approved for use in swine in Canada in 2004.

In pigs, florfenicol is typically delivered via intramuscular injection and is indicated for treatment of respiratory disease associated with bacteria such as *Actinobacillus pleuropneumoniae*, *Bordetella bronchiseptica*, *Glaesserella parasuis*, *Pasteurella multocida*, and *Streptococcus suis* and may be used off-label for other purposes. Intramuscular injection of florfenicol (15 mg/kg body weight) in pigs, with a plasma half-life of 11 to 18 h (2, 3) has been shown to result in concentrations of the drug in the lower gastrointestinal tract that are similar to or higher than that of orally dosed pigs (4). Resistance to florfenicol is often mediated through phenicol exporter genes such as *fexA*, *fexB*, or *floR* (1, 5) as well as the ribosomal protection protein genes *optrA* and *poxtA* and the 23S rRNA methyltransferase gene *cfr* (chloramphenicol-florfenicol resistance) (6). Significantly, many of the genes conferring resistance to florfenicol are carried on mobile genetic elements (MGEs) such as plasmids and transposons which enhance their dissemination among bacteria (7).

To date, there are no published studies that have investigated the effect of florfenicol use on the gut microbiome or resistome (all antimicrobial resistance genes) in pigs. Therefore, in this study, we administered florfenicol to piglets and collected fecal samples at several different time points until they reached market weight to determine the short- and long-term effects of florfenicol on the gut microbiome and resistome. We also collected colostrum, milk, vaginal, and fecal samples from their sows to assess the contribution of these sources to the piglet microbiome and resistome.

## METHODS

### Animals and experimental design

Pigs were cared for according to the guidelines of the Canadian Council on Animal Care. The Lacombe Research and Development Centre Animal Care Committee reviewed and approved all procedures and protocols involving all animals (animal use protocol number 201905). Landrace x Yorkshire sows (n =14) that had been inseminated with Duroc semen were given oxytocin to ensure that they all farrowed around the same time. Piglets were blocked by litter, weight, and sex and randomly assigned to the florfenicol (n = 30) or control groups (n = 30). At 1 and 7 days of age, piglets in the florfenicol treatment group were given an intramuscular injection of florfenicol (15 mg/kg body weight). Pigs from the two treatment groups had no contact with each other during the study and piglets consumed only sow milk prior to weaning at 21 days of age (i.e., no creep feed). Fecal swabs (FLOQSwabs; Copan, Murrieta, CA, USA) were collected from piglets at 1, 4, 7, 11, 14, 21, 28, 84, 112, and 140 days of age. Vaginal swabs were taken from the sows at farrowing (d 0) and fecal samples on d 0, 1, 4, 7, 11, 14, and 21. Colostrum was collected from the sows during or immediately after farrowing, and milk was collected on d 7 and 21. Approximately 30 minutes prior to the collection of milk, sows were injected with oxytocin to induce milk expression. The teat was thoroughly cleaned with 0.5% hydrogen peroxide prior to the collection of colostrum and milk into a sterile 150 ml screw cap container using sterile gloves. All samples were immediately placed on ice, transported to the laboratory, and stored at −80°C until DNA extraction. Pigs were also weighed on d 1, 7, 14, 21, 28, 70, 84, 112, and 140.

### Extraction of DNA

DNA was extracted from fecal and vaginal swabs using the QIAamp BiOstic bacteremia DNA kit (Qiagen, Mississauga, ON, Canada) as previously described (8). The DNeasy PowerFood Microbial kit (Qiagen) was used to extract DNA from the colostrum and milk samples. Fat was removed from the sample with a sterile cotton swab after centrifugation at 5,000 x g for 30 min at 4°C. The pellet was washed with 1.8 ml of phosphate-buffered saline (PBS), centrifuged at 10,000 x g, and the remaining fat layer removed. The wash step was repeated three to five times until all the visible fat was removed. The pellet was subjected to enzymatic lysis with 90 µl of 50 mg/ml lysozyme (Roche, Mannheim, Germany) and 50 µl of 5 KU/ml mutanolysin from *Streptomyces globisporus* ATCC 21553 (MilliporeSigma, Oakville, ON, Canada) at 55°C for 15 min followed by the addition of 28 µl of 20 mg/ml proteinase K (MilliporeSigma) for an additional 15 min at 55°C. The suspension was then centrifuged at 10,000 x g for 1 min at room temperature and the supernatant discarded. The pellet was suspended in 450 µl MBL buffer (Qiagen) and transferred to a PowerBead tube (Qiagen). The PowerBead tube was vortexed for 5 s and then placed into a thermoshaker set at 65°C and 1,400 rpm for 10 min. Following incubation, the samples were bead-beaten in a MP FastPrep-24 (MP Biomedicals, Solon, OH, USA) at 6 m/s for 90 s. After a 5 min rest, the samples were centrifuged at 13,000 x g for 1 min at room temperature. The supernatant was then transferred to a clean collection tube and the manufacturer’s protocol was followed for all subsequent steps.

### 16S rRNA gene sequencing and analysis

The V4 region of the 16S rRNA gene in the extracted DNA from all fecal, colostrum/milk, and vaginal samples was amplified and sequenced as previously described (9). Cutadapt v. 3.4 (10) was used to remove primer sequences and any reads shorter than 215 bp. The reads were then processed using DADA2 v. 1.20.0 (11) in R v. 4.1.0 with forward and reverse reads trimmed to 200 bp each and merged with a minimum overlap of 75 bp. Amplicon sequence variants (ASVs) were resolved, chimeras removed, and taxonomy assigned to the ASVs using the SILVA SSU database release 138.1 (12). ASVs classified as chloroplasts, mitochondria, or eukaryota were removed. Extraction kit controls (n =18) were also included; however, only one ASV classified as *Escherichia coli* was identified in these negative controls and at a significantly lower abundance than in the biological samples and was therefore retained. A 20-strain whole-cell mock community was also extracted and sequenced (MSA-2002; ATCC). The pig fecal samples were randomly subsampled to 5,200 sequences prior to the calculation of ASV richness and the inverse Simpson diversity as well as Bray-Curtis dissimilarities using Phyloseq v. 1.36.0 (13) and vegan 2.6-4 (14).

### Shotgun metagenomic sequencing and analysis

Fecal samples from a random subset of 16 pigs per treatment group collected on d 1, 4, 7, 11, 14, 21, 28, 84, and 140 were selected for shotgun metagenomic sequencing. The sow fecal samples from d 1, 7, and 21 as well as all colostrum/milk and vaginal swabs collected were also included. Metagenomic libraries were prepared and sequenced on a NovaSeq 6000 instrument with a S4 flow cell (300 cycles) (Illumina Inc, San Diego, CA, USA) as previously detailed (15). Low-quality reads and sequencing adapters were removed with fastp v.0.23.2 (16) using a 4-bp sliding window and a quality threshold of 15. Reads shorter than 100 bp were also removed. To remove swine host and PhiX sequences, the reads were aligned to three swine genome assemblies (Sscrofa11.1 [Duroc], Berkshire_v1 [Berkshire], USMARCv1.0 [Duroc x Landrace x Yorkshire]) and the *Escherichia* phage phiX174 genome (NC_001422) with Bowtie2 v.2.4.4 (17). Reads that did not map to these genomes were then extracted using SAMtools v.1.14 (18) and BEDtools v.2.30.0 (19). The metagenomic reads were taxonomically classified using Kraken2 v. 2.1.3 (20) and Bracken v. 2.9 (21) with the Genome Taxonomy Database (GTDB) release 220 (22). The resistance gene identifier (RGI) v. 6.0.2 and the Comprehensive Antibiotic Resistance Database (CARD) v.3.2.6 (23) with KMA (24) were used to screen the unassembled reads for antimicrobial resistance genes (ARGs).

### Metagenome-assembled genomes

The metagenomic reads were also individually assembled for each sample as well as co-assembled depending on the sample type and treatment group using MEGAHIT v. 1.2.9 (25). Briefly, colostrum and milk, vaginal, and sow fecal samples were all co-assembled separately. Fecal samples from the control and florfenicol-treated pigs before and after weaning were also co-assembled separately. Each sample was mapped back to its respective assembly and co-assembly using Bowtie2 and the contigs (≥ 2000 bp) were binned into metagenome-assembled genomes (MAGs) using MetaBAT 2 v. 2.2.15 (26). The completeness and contamination of the co-assembled and individually assembled MAGs were calculated with CheckM v1.2.0 (27) resulting in 78,968 MAGs that were at least 90% complete and had less than 5% contamination. These MAGs were then dereplicated using dRep v.3.2.2 with primary clustering set at 90% average nucleotide identity (ANI) and secondary clustering set at 99% ANI. Taxonomy was then assigned to the 1,546 dereplicated MAGs using GTDB-Tk v. 2.4.0 (28) and the GTDB release 220.

Antimicrobial resistance genes were identified in the MAGs using the RGI with the CARD. The MAGs were also screened for virulence genes with DIAMOND v. 2.0.15.153 (29) and the virulence factor database (VFDB) v. 20230915 (30) at 90% identity. *E*. *coli* MAGs were classified by serotype using SerotypeFinder v.2.0 (31). Multilocus sequence typing (MLST) was done on MAGs assigned to a species that was in the PubMLST database (32). PhyloPhlAn v. 3.0.67 (33) was used to create a phylogenomic tree of the MAGs by aligning 400 universal marker genes. The relative abundance of each MAG in each sample was calculated using CoverM v. 0.6.1 (https://github.com/wwood/CoverM).

### Quantification of *floR*

As *floR* was the most differentially abundant ARG between the treated and untreated pigs, the absolute concentration of this gene was determined using qPCR. Briefly, the primers *floR*-F 5’-CGGTCGGTATTGTCTTCACG-3’ and *floR*-R 5’-TCACGGGCCACGCTGTAT-3’ (34) were used to target and amplify the *floR* gene in the same samples previously selected for shotgun metagenomic sequencing. Each reaction contained the forward (7.5 pmol) and reverse primers (5 pmol), 25 to 50 ng DNA template, and Brilliant II SYBR Green qPCR Master Mix (Agilent Technologies, Mississauga, ON, Canada) in a total volume of 20 µl. The three-step PCR consisted of an initial denaturation at 95°C for 10 min followed by 40 cycles of annealing at 54°C for 1 min and extension at 72°C for 30 s as per manufacturer’s instructions. Fluorescence was detected during annealing and extension on an AriaMx real-time PCR system (Agilent Technologies). A dissociation curve was generated at the end of each qPCR run with an increase in temperature of 0.5°C/s up to 95°C to ensure that only one amplicon was produced. A negative control and a serially diluted seven-point standard curve from 1.42 x 10^2^ to 1.42 × 10^8^ copies were included with each qPCR run. The standard consisted of the *floR* gene in a pUCIDT (Amp) vector (Integrated DNA Technologies, Coralville, IA, USA). The DNA concentration of the *floR* standard was determined with a dsDNA HS assay kit (Thermo Fisher Scientific). Copy number was calculated using the stock *floR* standard concentration, the plasmid size, and a molar mass of 650 g/mol per bp. All samples and standards were analyzed in triplicate.

### Culturing and sequencing of florfenicol-resistant bacteria

Florfenicol-resistant isolates were recovered from fecal swabs (n = 11) from florfenicol-treated piglets on days 4 and 11 to evaluate the genomic context of certain florfenicol resistance genes. Briefly, swabs were added to 10 ml of brain heart infusion (BHI) broth (Oxoid, Basingstoke, Hampshire, UK) in Hungate anaerobic tubes (Chemglass Life Sciences, Vineland, NJ, USA), and incubated for 4 h at 39°C. Serial dilutions were then plated onto BHI agar (Oxoid) containing 32 µg/ml florfenicol (Thermo Fisher Scientific, Waltham, MA, USA) and incubated for 24 h at 39°C under anaerobic conditions. One colony was selected from each agar plate, re-streaked onto BHI agar with 32 µg/ml florfenicol, and incubated at 39°C for 16 h. From these plates, one colony was inoculated into 10 ml of BHI broth in a Hungate tube and incubated for 6 h at 39°C. These cultures were then stored in BHI with 20% (v/v) glycerol at −80°C until DNA extraction.

A DNeasy Blood & Tissue Kit (Qiagen) was used to extract DNA from each culture as per manufacturer’s protocol for Illumina sequencing. The DNA concentration was determined using a Qubit dsDNA HS assay kit (Thermo Fisher Scientific) and 500 ng of DNA was used with an Illumina DNA Prep kit (Illumina Inc.) to generate genome sequence libraries as described by the manufacturer. The size of each sample library was checked using a D1000 ScreenTape System (Agilent Technologies) as per manufacturer’s instructions, and the concentration of each library was analyzed with a Qubit dsDNA HS assay kit. Libraries were normalized to 4 nM prior to pooling and denaturing. The libraries were then diluted to 11 pM and 1% denatured PhiX (Illumina Inc.) was added as described by the supplier. The genomic libraries were then sequenced on a MiSeq instrument with a MiSeq Reagent Kit v2 (300 cycles; 2 x 150 bp) (Illumina Inc.).

The florfenicol-resistant isolates were also sequenced using a MinION Mk1B device with R10.4.1 flow cell (Oxford Nanopore Technologies, Oxford, UK). Briefly, high molecular weight (HMW) DNA was extracted from an overnight culture using the Monarch HMW DNA Extraction Kit for Tissue (New England Biolabs, Ipswich, MA, USA) as per manufacturer’s protocol for HMW DNA extracted from bacteria. DNA concentration was determined with a Qubit dsDNA HS assay kit and the size and quality of the DNA were evaluated with a genomic DNA ScreenTape system as per manufacturer’s instructions. Library preparation was done with the Native Barcoding Kit 24 V14 following the protocol supplied by the manufacturer. HMW DNA from *E*. *coli* was difficult to elute from the AMPure XP Beads after the DNA repair and end-prep step; therefore, the input DNA concentration was increased to 1 µg from 400 ng, the volume of nuclease-free water to solubilize the HMW DNA was increased to 20 µl from 10 µl and the incubation time was increased to 10 min from 2 min.

During the adapter ligation and cleanup step, the long fragment buffer supplied in the kit was used to enrich for DNA fragments over 3 kb in length. For all flow cells the recommended amount of final prepared library was loaded with the library beads supplied in the kit. The flow cell was primed with the recommended addition of ultrapure bovine serum albumin (Invitrogen, Waltham, MA, USA) to a final concentration of 0.2 mg/ml. During data acquisition and base calling, the default parameters were used in the MinKNOW v. 23.11.4 software installed with Bream v.7.8.2, Dorado 7.2.13 and MinKNOW core v. 5.8.6. After the sequencing run was completed, base calling was re-done with the high accuracy model within MinKNOW.

### Genomic analysis of florfenicol-resistant isolates

Genomes were assembled using the MinION reads and polished with Illumina reads as per Wick et al. (35). Briefly, all MinION reads were filtered with Filtlong v. 0.2.1 (https://github.com/rrwick/Filtlong) removing reads shorter than 6,000 bp and retaining only the best 90% of reads. Filtered reads for each sample were then subsampled into 12 read subsets with Trycycler v. 0.5.4 (36) and an assembly was then made with each read subset. The four assemblers used were: Flye v. 2.9.2-b1786 (37) with the nano-hq flag, miniasm v. 0.3-r179 (38) with minimap2 v. 2.26-r1175 (39) and polishing by minipolish v. 0.1.2 (40), Raven v. 1.8.3 (41) with Racon v. 1.5.0 (42) and the disable-checkpoints flag, and Canu v. 2.2 (43). All contigs generated from each of the assemblers for each sample were then grouped into clusters using Trycycler. Clusters were discarded if most contigs in the cluster did not have a similar size and depth. “Trycycler reconcile” was run on each cluster with contigs and clusters removed if they could not be reconciled.

A multiple sequence alignment was made from each reconciled cluster with “Trycycler msa”, and then each read was assigned to a cluster using “Trycycler partition”. A consensus sequence was then generated from each cluster with “Trycycler consensus”. The consensus sequence for each cluster was polished with Medaka v. 1.11.1 (https://github.com/nanoporetech/medaka) using the r1041_e82_400bps_hac_v4.2.0 model. The polished assemblies were concatenated into one assembly per sample. Prior to polishing the long-read assemblies with Polypolish v. 0.5.0 (44), the Illumina reads were filtered with fastp, aligned to the polished assembly for each sample with bwa v. 0.7.17-11 (45), indexed with bwa, and size filtered with Polypolish. Taxonomy was assigned to the polished genome assemblies with GTDB-Tk and the GTDB release 220. Assembly metrics were assessed with QUAST v. 5.2.0 (12) and contamination and completeness were determined with CheckM2 v. 1.0.1 (46). The finished genomes were screened for virulence genes and ARGs and the *E*. *coli* isolates were assigned to serotypes as described above for the MAGs. Plasmids from the polished genomes were classified based on their replicon type using PlasmidFinder v. 2.1.6-1 (47). The ANI between genomes was calculated with fastANI v. 1.33 (48) and the isolate genomes and their associated plasmids were viewed with Proksee (49).

The susceptibility of the *Enterococcus* spp. isolates to linezolid was also assessed. Briefly, linezolid (MilliporeSigma) was dissolved in molecular grade water to a concentration of 2.5 mg/ml and then diluted in cation-adjusted Mueller-Hinton (MH) Broth 2 (MilliporeSigma) to a concentration of 128 µg/ml. This linezolid working stock was serially diluted two-fold into designated wells in a 96-well microplate (Thermo Scientific). To prepare the inoculum for each isolate, the culture was streaked onto BHI agar and grown overnight (16 h) at 39°C. Colonies from the plate were then added to MH broth until the suspension resembled a 0.5 McFarland standard and 100 µl of this suspension was added to 9.9 ml of MH broth. Finally, 50 µl of this inoculum was added to the designated wells. The inoculated microplate was grown for 48 h at 35°C and the lowest linezolid concentration without visible growth was determined to be the minimum inhibitory concentration (MIC). The susceptibility assay was performed in duplicate on three separate days. Two enterococci strains known not to be carrying any oxazolidinone resistance genes, *Enterococcus faecium* ATCC 35667 and *Enterococcus faecalis* ATCC 29212, were included as controls. Interpretation of the MICs was based on the Clinical and Laboratory Standards Institute (CLSI) breakpoints of ≤ 2 µg/ml = sensitive, 4 µg/ml = intermediate resistance, and ≥ 8 µg/ml = resistant (CLSI, Standards for Antimicrobial Susceptibility Testing, M100).

### Statistical analysis

Differentially abundant microbial species, ARGs, and MAGs between the florfenicol-treated and control pigs were identified with MaAsLin2 v. 1.16.0 (50) in R 4.3.2. Only those microbial species with a percent relative abundance of at least 0.1% and ARGs present in at least 25% of the samples being analyzed were included. Permutational multivariate analysis of variance (PERMANOVA) of the Bray-Curtis dissimilarities was calculated in vegan to determine the effect of florfenicol treatment on the structure of the microbial community (ASVs from 16S rRNA gene data) and resistome (ARGs). The effect of florfenicol on average daily gain was evaluated using a linear mixed model with individual pig as the random effect and treatment and age as the fixed effects with the R package lme4 v. 1.1-32 (51). Post-hoc comparisons were carried out within each time period using Tukey’s honestly significant difference. Sourcetracker2 v. 2.0.1-0 (52) was used to predict the relative contribution of the sow vagina, colostrum, milk, and feces to the piglet gut microbiome. For this analysis, the samples included were limited to those from the control piglets and their sows to avoid any treatment effect. The 16S rRNA gene sequences randomly subsampled to 2,000 sequence per sample were used to assess the contributions to the microbiome while the antimicrobial resistance gene data from the RGI was used for the resistome assessment.

## RESULTS

### Metagenomic sequencing summary

After quality filtering and host DNA removal, 1.85 Tb of sequence data from 368 metagenomic samples were available for downstream analyses. As expected, given the difficulty in removing host cells from colostrum and milk samples prior to DNA extraction, these samples had a very low microbial to host DNA ratio (up to 99% host DNA) and vaginal swabs were also heavily contaminated with DNA from the sow (Table S1). The metagenomes of the mock community were also assembled and binned into MAGs. All 20 bacterial species in the mock community were represented among these MAGs, although with varying levels of completeness, and no species not included in the mock community were identified (Table S2).

### Animal performance

The average daily gain differed significantly between the control and florfenicol-treated pigs only during the 1 to 7 d period (Fig. S1). Notably, it was the florfenicol-treated piglets that grew slower during this time frame. As expected, the average daily gain for piglets was reduced during the one-week post-weaning period as they adjusted to solid feed.

### Effect of florfenicol on the piglet gut microbiome

Florfenicol administration had the greatest effect on the piglet fecal microbial community structure on day 4, three days after the first treatment, based on the 16S rRNA gene sequence data (PERMANOVA: R^2^ = 0.18; P = 0.0003; Fig. 1). A second injection on day 7 increased the dissimilarity of the microbiomes of the two groups of pigs; however, this effect was lost by day 21, just prior to weaning. Although the effect was relatively small, the control and florfenicol-treated pigs did have significantly different microbiomes on days 28, 84, and 140, suggesting there may have been some minor residual effects of florfenicol treatment post-weaning. The florfenicol-treated piglets had a less rich (fewer ASVs) microbiota on days 4, 11, and 28, although the opposite was true at the end of the study (d 140) (Fig. S2). In terms of microbial diversity, the control piglets had greater microbial diversity (inverse Simpson diversity index) than the florfenicol group only on d 28 (Fig. S2).

**Figure 1.**
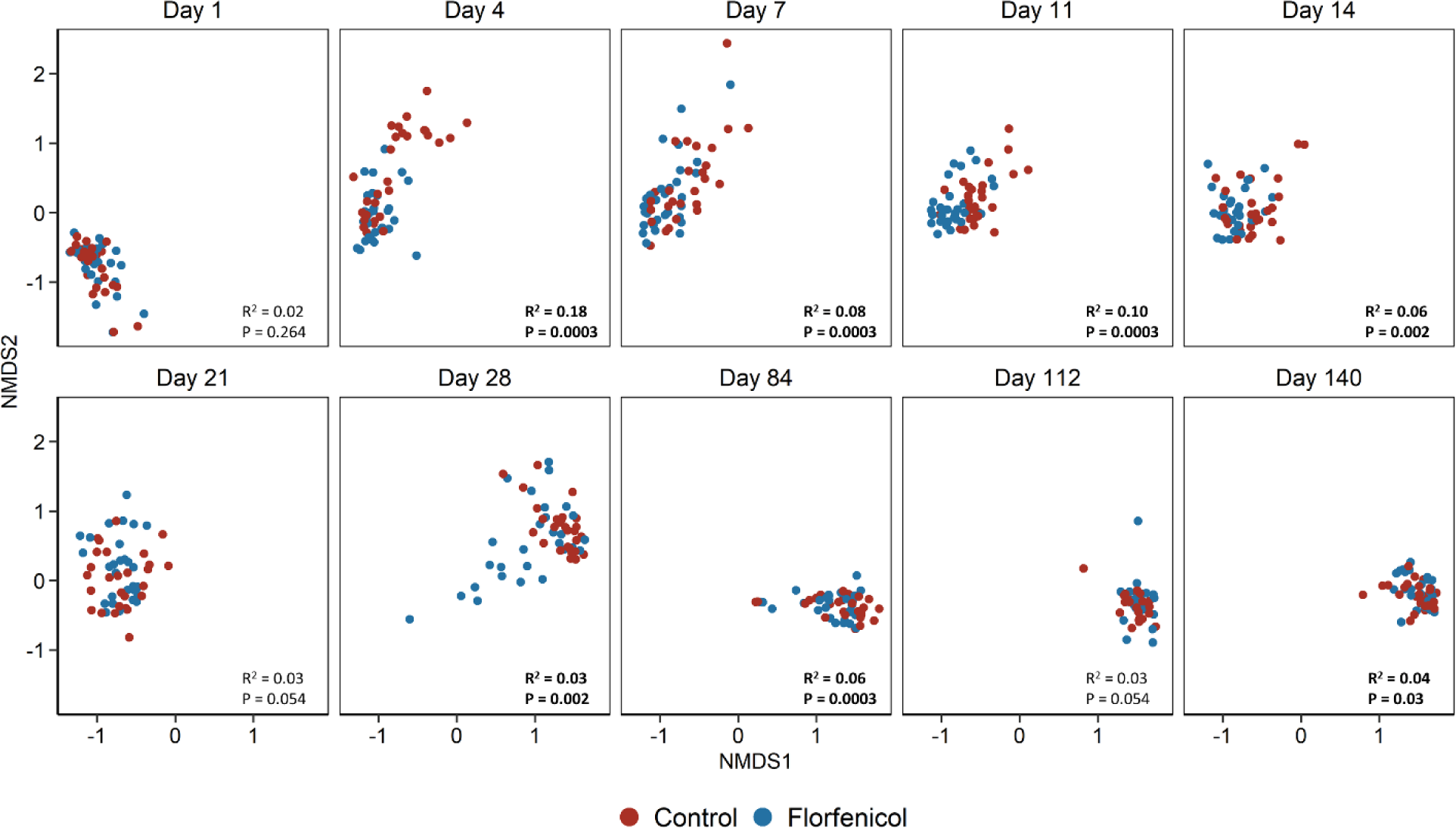
Non-metric multidimensional scaling (NMDS) plots of the Bray-Curtis dissimilarities for untreated pigs (control; n = 30) and pigs treated with florfenicol (n = 30) on days 1 and 7 based on 16S rRNA gene sequences. Permutational multivariate analysis of variance (PERMANOVA) R^2^ and P-values are included in each plot.

Based on the analysis of the metagenomic sequences, there were 67 bacterial species that were differentially abundant between the control and florfenicol-treated pigs on d 4 (Fig. 2; Table S3). *Fusobacterium mortiferum*, *Ruminococcus gnavus*, and *Pauljensenia hyovaginalis* were among the species strongly associated with the control pigs at this time while *Anaerotignum* sp001304995, *Escherichia ruysiae*, and *E. faecalis* were among the 28 bacterial species relatively more abundant in pigs treated with florfenicol. There were progressively fewer bacterial species that were differentially abundant between the two groups of pigs from d 7 through 21, just prior to weaning. Nine bacterial species were consistently differentially abundant between the control and florfenicol-treated pigs on d 4, 7, and 11 (Fig. 3). Among these species, *Basfia rossii* (NCBI: *Actinobacillus rossii*) was also enriched in the gut microbiomes of the control pigs from days 4 through 21 while CALYPF01 sp945271245 and *Clostridium scindens* were associated with the florfenicol-treated pigs during this period. Although no bacterial species were differentially abundant between the two groups at one-week post-weaning (d 28) and at the end of the study (d 140), 12 species were differentially abundant on d 84 (Table S3). There was also one species, *Holdemanella porci*, that was significantly associated with the control pigs on d 84 in addition to days 4, 7, and 11.

**Figure 2.**
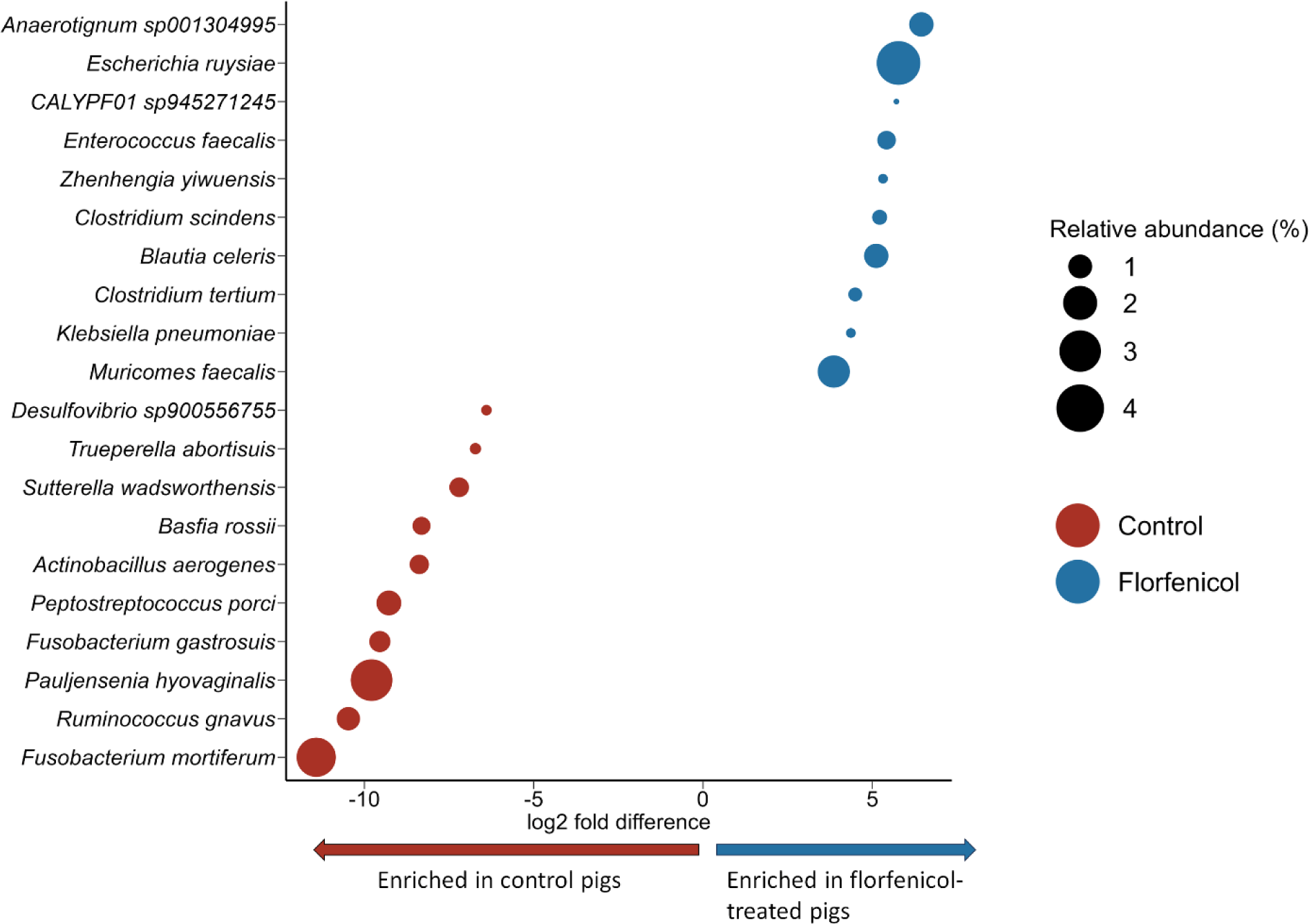
The 20 most differentially abundant bacterial species between control (n = 16) and florfenicol-treated (n = 16) piglets at 4 days of age. The size of the circles is proportional to the relative abundance abundance (%).

**Figure 3.**
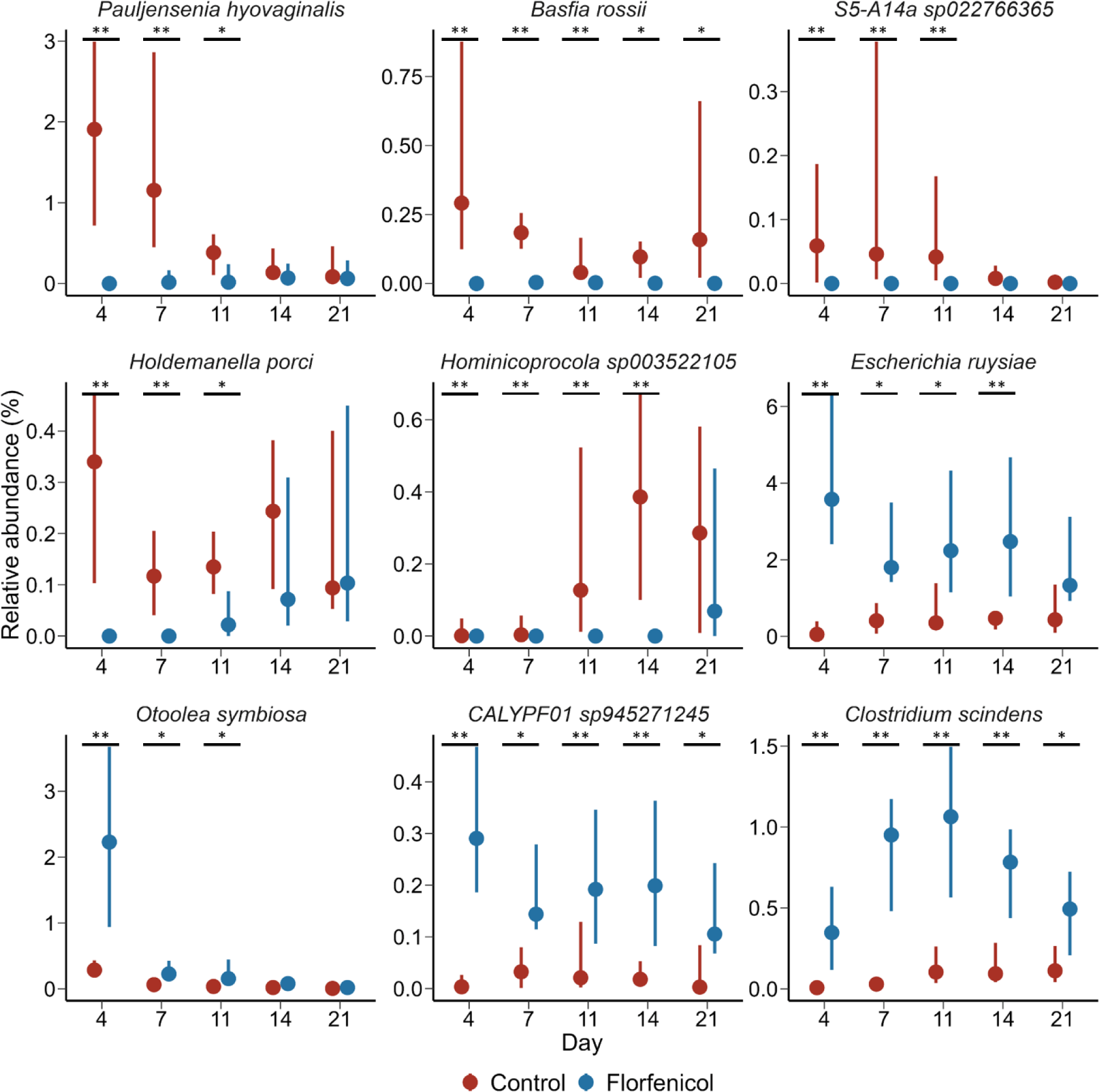
Percent relative abundance of bacterial species that were consistently differentially abundant between control (n = 16) and florfenicol-treated (n = 16) piglets on days 4, 7, and 11 (n = 16). The point represents the median value and the lines are the 25^th^ and 75^th^ percentiles. * = P-value < 0.05; ** = P-value < 0.01.

### Effect of florfenicol on the piglet gut resistome

Similar to its effect on the piglet gut microbiome, florfenicol had the greatest effect on the structure of the resistome on day 4 (Fig. S3; R^2^ = 0.25; P < 0.05). As may be expected, ARGs conferring resistance to the phenicols were relatively more abundant in the pigs treated with florfenicol from days 4 through 28 (Fig. 4; P < 0.05). However, there was also evidence that florfenicol was co-selecting for resistance to other antimicrobial classes as ARGs that confer resistance to aminoglycosides, beta-lactams, peptides, and sulfonamides were enriched in the metagenomes of treated piglets during at least one sampling period between 4 and 21 days of age. Conversely, ARGs conferring resistance to the macrolides-lincosamides-streptogramin B (MLS_B_) class were relatively more abundant in the control piglets on d 4 and 7, although the opposite was observed at d 84. The abundance of MLS_B_ and tetracycline ARGs in the pig gut remained relatively consistent from d 1 through 140, similar to the levels observed in the sows (Fig. 4).

**Figure 4.**
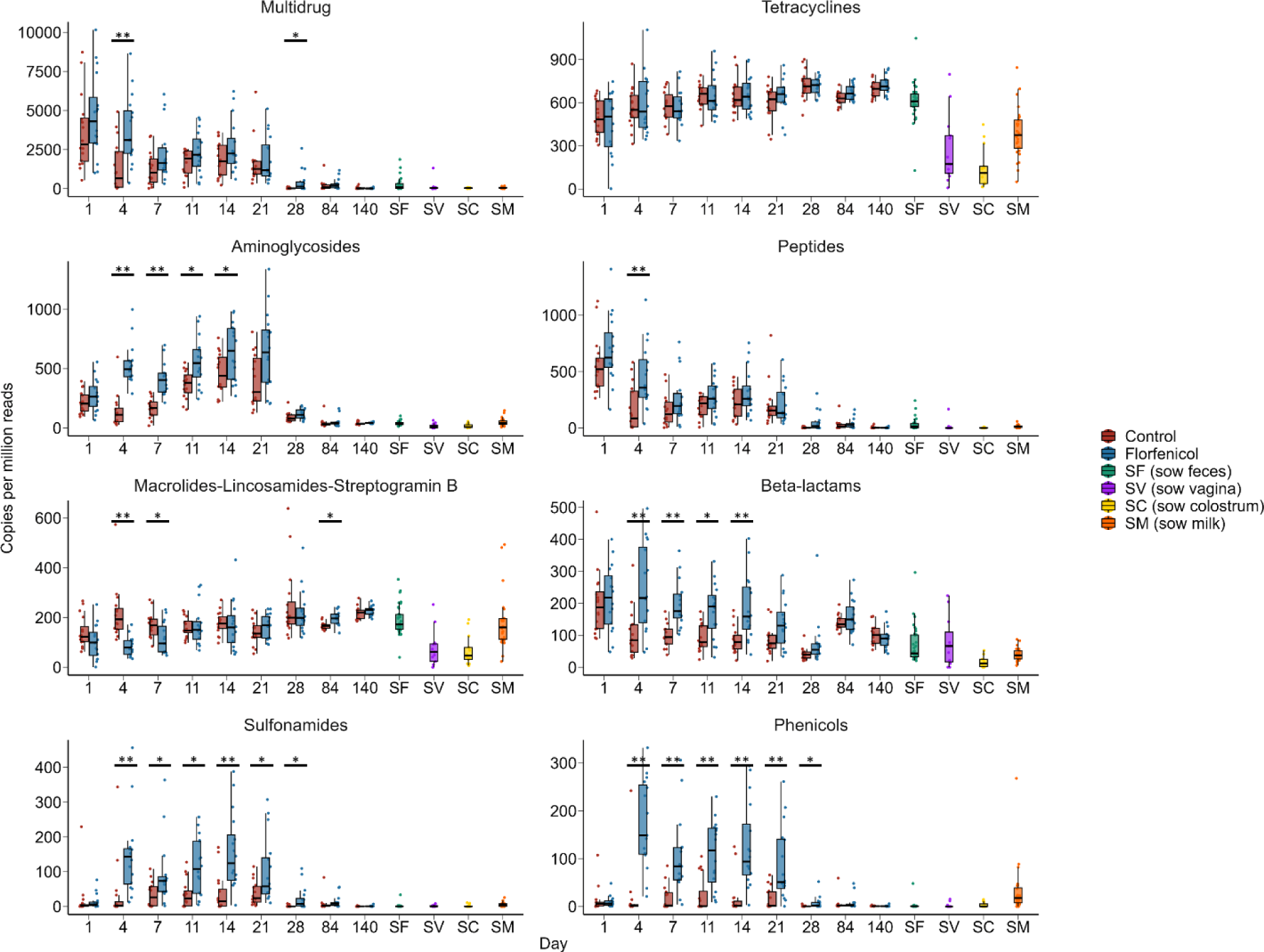
The relative abundance of antimicrobial resistance genes by antimicrobial class they confer resistance in the metagenomes of control (n = 16) and florfenicol-treated pigs (n = 16). Also included are sow fecal (SF), vaginal (SV), colostrum (SC), and milk (SM) samples. * P-value < 0.05; ** P-value <0.01.

In terms of individual ARGs, 111 were differentially abundant in the piglet fecal metagenomes on day 4, of which 91 were enriched in the florfenicol-treated group (Table S4). The *floR* and *fexB* genes, which confer resistance to florfenicol, were most strongly associated with the florfenicol-treated piglets. These two ARGs were also relatively more abundant in the feces of the florfenicol-treated piglets from days 4 through 21. We also quantified the *floR* gene using qPCR and the results largely mirrored that of the metagenomic data with the exceptions of days 7 and 21 (Fig. 5). The means of the two groups were clearly separated on these days, however, insufficient DNA was available for many of the samples from these two time points after 16S rRNA gene and metagenomic sequencing and therefore statistical power was reduced. Another florfenicol-specific resistance gene, *fexA*, was also enriched in the piglets treated with florfenicol on days 4 through 14. The *fexA*, *fexB*, and *floR* genes all encode for phenicol-specific efflux pumps. The ARGs *cfr*, *cfr*(B), and *clc*D encode 23S ribosomal RNA methyltransferases that confer resistance to phenicols along with other antimicrobial classes such as MLS_B,_ oxazolidinones, and pleuromutilins (53). Similarly, resistance to phenicols, as well as oxazolidinones, can be mediated through the ABC-F subfamily ATP-binding ribosomal protection protein genes *optrA* and *poxtA* (54). All five of these ARGs (*cfr*, *cfr*(B), *clcD*, *optrA*, *poxtA*) were relatively more abundant in the gut microbiomes of florfenicol-treated piglets during at least one sampling time from d 4 through d 21. In addition to *fexB*, *floR*, and *clcD*, *bla*_CMY-59,_ a class C beta-lactamase gene, and *tet*(A), a tetracycline efflux pump gene, were also consistently relatively more abundant in the piglets administered florfenicol on d 4 through d 21. With the exception of *tet*(A) and *tet*(D), the relative abundance of several *tet* genes, namely *tet*(B), *tet*(O), *tet*(O/M/O), *tet*(Q), *tet*(T), *tet*(O/M/O), *tet*(W/32/O), *tet*(X1), and *tet*(Z), was higher in the control piglets on at least one sampling time through d 4 to d 21 (Table S4). Similarly, certain MLS_B_ resistance genes including *erm*(33), *erm*(47), *erm*(X), *lnu*(C), *lnu*(P), *lsa*(C), and *vat(*E) were relatively more abundant in the control piglets.

**Figure 5.**
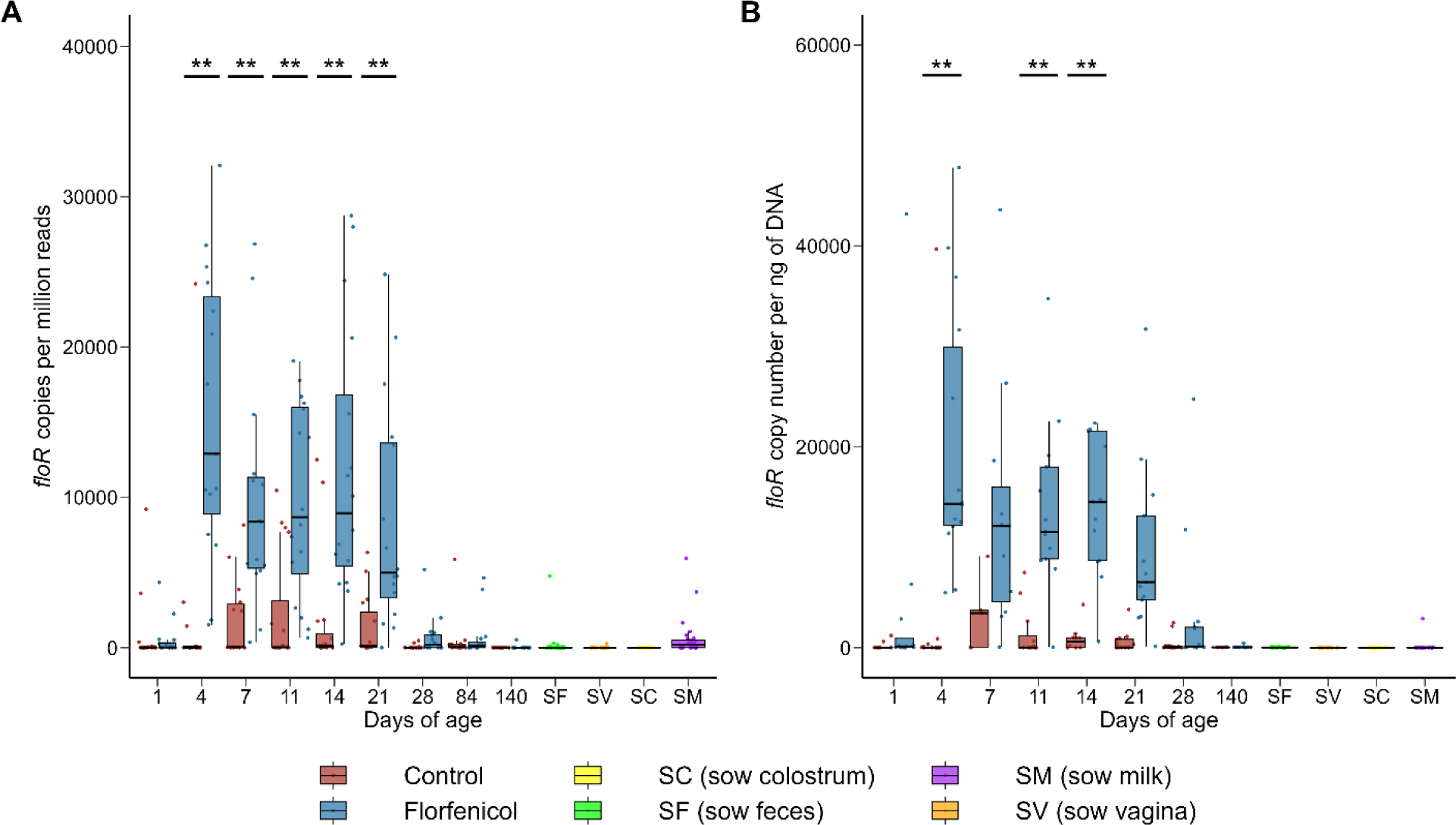
The relative abundance (A) and concentration (B) of the *floR* gene determined by metagenomic sequencing and qPCR, respectively, in the feces of control (n = 16) and florfenicol-treated (n = 16) pigs. Also included are sow fecal (SF), vaginal (SV), colostrum (SC), and milk (SM) samples. * P-value < 0.05; ** P-value <0.01.

### Sow colostrum, milk, vaginal, and fecal microbiomes and resistomes

The sow colostrum, milk, vaginal, and fecal microbiomes were also characterized to assess the potential contributions of these sources to the piglet gut microbiome and resistome. Based on metagenomic sequencing, the relatively most abundant archaeal/bacterial species in the colostrum were *Lactobacillus amylovorus*, *Methanobrevibacter* sp900769095, *Streptococcus thoraltensis*, *Prevotella* sp945863825, *Rothia nasimurium*, *Lelliottia chinensis*, and *Cutibacterium acnes* (> 2% relative abundance; Table S5). In milk samples collected on days 7 and 21, the bacterial species *Rothia nasimurium* was relatively most abundant. Similar to the colostrum, the potentially beneficial bacterial species *L. amylovorus* was among the relatively most abundant microbes in the milk samples (≥ 2.6%). However, unlike the colostrum, *S*. *suis*, a known pathogen in pigs, was relatively abundant in the milk samples at both sampling times (≥ 2.8%). The vaginal microbiome of the sows during farrowing was largely dominated by *B*. *rossii* (30.6 ± 8.0% SEM), *Streptococcus thoraltensis* (7.2 ± 3.1%), *Veillonella caviae* (3.8 ± 1.2%), as well as *S*. *suis* (4.2 ± 2.7%). *Bacteroides fragilis* was relatively most abundant in the feces of sows during the nursing period along with *Arcanobacterium* sp028724905, *E*. *coli* (particularly on d 0), *Mobiluncus porci*, and *Prevotella faecis* (Table S5).

Using the 16S rRNA gene sequences of the control pigs and their sows, colostrum was identified as a major contributor to the piglet gut microbiome on day 1 in particular as well as up to day 28 (Fig. 6A). Sow feces were also predicted to be a significant source for the piglet gut microbiome from day 4 through 140, although there was considerable variation especially in the nursing period. For the piglet resistome, sow feces were predicted to be the greatest source of ARGs throughout the entire production cycle (Fig. 6B). Although highly variable between samples, the colostrum, milk, and vagina carried ARGs conferring resistance to the MLS_B_, tetracycline, and beta-lactam classes (Figure 4).

**Figure 6.**
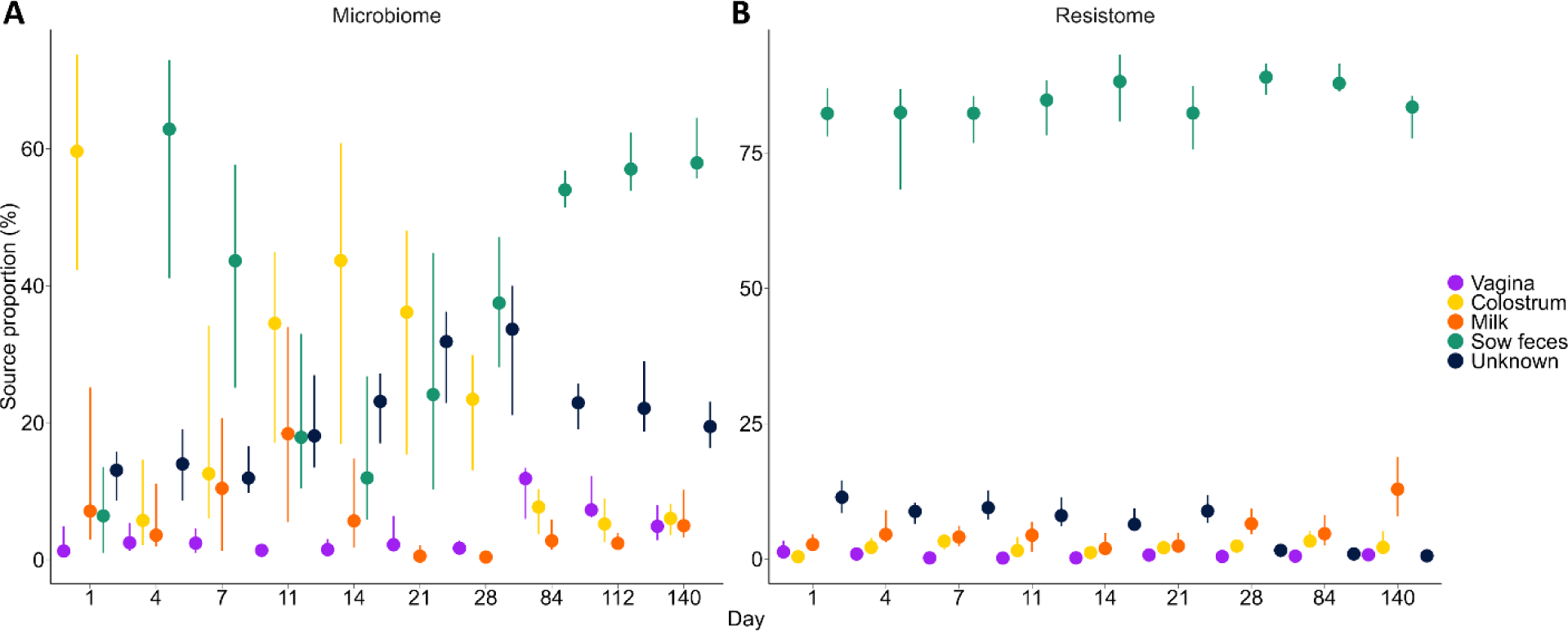
Predicted contributions of the vagina, colostrum, milk, and feces of the sows to the piglet gut microbiome (A; 16S rRNA gene sequences; n = 30) and resistome (B; metagenomes; n = 16). The points represent median values and the lines are the 25^th^ and 75^th^ percentiles.

### Metagenome-assembled genomes

A total of 1,546 non-redundant MAGs with less than 5% contamination and greater than 90% completeness were recovered from the piglet fecal and the sow colostrum, milk, fecal, and vaginal samples (Fig. 7; Table S6). The most frequently identified species represented by these MAGs were all uncultured species: *Sodaliphilus* sp004557565 (62 MAGs), *Collinsella* sp002391315 (29 MAGs), *Campylobacter* sp945873855 (24 MAGs), *Prevotella* sp000434975 (22 MAGs), and CAG-349 sp003539515 (Christensenellales; 19 MAGs). There were also 19 *Escherichia coli* MAGs. The vast majority of reads from the colostrum and milk samples did not align to any of the MAGs (91.8% and 78.0%, respectively); however, among those that did, *L. amylovorus* and *Rothia* spp. MAGs were proportionally most abundant (Table S6). Similar to the unassembled reads, MAGs assigned to *B*. *rossii*, *S*. *thoraltensis*, *S*. *suis, Phocaeicola* sp022764505, and *V*. *caviae* had the greatest relative abundance (> 0.5%) in the sow vaginal microbiome at farrowing. Overall, *B. fragilis*, *Sarcina* (*Clostridium*) *perfringens*, *E*. *coli*, *Lactobacillus delbrueckii*, and *Limousia pullorum* MAGs were relatively most abundant in pig fecal samples, although there was considerable variation by sampling time.

**Figure 7.**
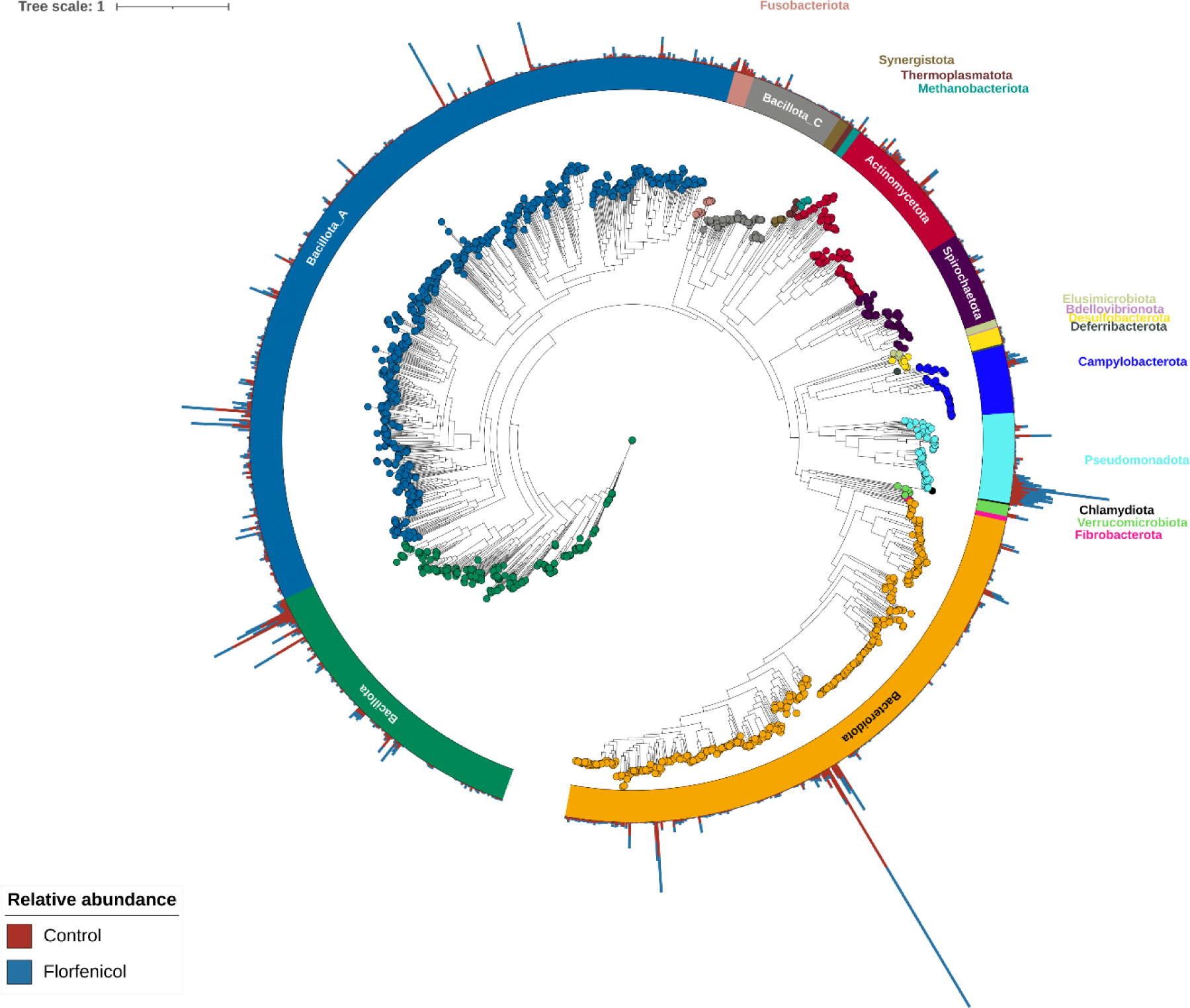
Maximum likelihood phylogenetic tree of the metagenome-assembled genomes (MAGs) recovered from pig fecal and sow colostrum, fecal, milk, vaginal metagenomes based on the alignment of 400 marker genes. MAGs are colored and labeled by GTBD-tk assigned phyla. The outer bars display the overall percent relative abundance (0% to 3.6%) of each MAG in the fecal microbiomes of control vs. florfenicol-treated pigs.

On day 4, a total of 209 MAGs were differentially abundant in the fecal microbiomes of the control (156 MAGs) and florfenicol-treated (53 MAGs) piglets (Table S7). Among the 53 MAGs that were relatively more abundant in the florfenicol-treated piglets were the 19 *E*. *coli* MAGs as well as MAGs classified as *Eisenbergiella porci*, *E*. *faecalis,* and *E*. *faecium*. All 29 *Collinsella* sp002391315 MAGs in the dataset were relatively more abundant in the feces of the untreated piglets at this time as were 14 *Prevotella* sp000434975, 8 *P*. *hyovaginalis*, 7 *Desulfovibrio* sp900556755, and 7 *Peptostreptococcus porci* MAGs. Other notable MAGs enriched in the control piglet fecal microbiomes on d 4 were *Campylobacter coli*, *Clostridioides difficile*, *Streptococcus hyovaginalis*, and *S*. *suis*. There were 55 differentially abundant MAGs on d 7, 87 MAGs on d 11, 58 MAGs on d 14, 2 MAGs on d 21, and none on d 28. Only one MAG, enriched in the control pig fecal samples and classified as *Peptoniphilus* sp022767005 (SUG3067), was consistently differentially abundant between the two groups of piglets from d 4 through d 21. Interestingly, on d 84 there were 47 differentially abundant MAGs between the two groups of pigs, with 38 of these MAGs more relatively abundant in the control pigs. The majority of these MAGs (27) were classified as *Collinsella* sp002391315. At the next sampling time, d 140, only 8 MAGs differed in relative abundance, and none were shared between days 84 and 140.

The MAGs were also screened for ARGs with 538 MAGs found to be carrying at least one ARG (Table S8). As expected, the 19 *E*. *coli* MAGs encoded the greatest number of ARGs as many of these genes are intrinsic to this species. The most widely encoded ARGs were *vanG* (206 MAGs), which typically confers only low-level resistance to vancomycin (55), and *adeF* (97 MAGs), a multidrug efflux pump gene. Although neither *fexB* nor *floR* were identified in any of the MAGs, *fexA* was binned into a *Staphylococcus borealis* MAG and *ant(4’)-Ib, fexA,* and *optrA* were found in a MAG classified as *Vagococcus lutrae.* The *poxtA* gene was detected in 38 MAGs, 7 of which were relatively more abundant in the florfenicol-treated pigs during at least one sampling time. These 7 MAGs included those assigned to *Eisenbergiella porci*, *Enterocloster clostridioformis*, *Enterocloster porci*, and *Hungatella hathewayi*. Genes conferring resistance to tetracyclines (*tet* genes) were identified in 152 different MAGs with *tet*(Q) (49 MAGs), *tet*(T) (36 MAGs), and *tet*(36) (35 MAGs) most prevalent. In addition, a *P. porci* MAG carried *cfr*(B) along with *lnu*(P) and a MAG designated *Clostridium* sp900759995 encoded *cfr*(B), *lnu*(C), and *tet*(36). *Clostridium* sp900759995 was relatively more abundant in the metagenomes of florfenicol-treated piglets on day 4, although the *P*. *porci* MAG was enriched in the untreated piglets on days 4 and 7.

The 19 *E*. *coli* MAGs were further classified using *in silico* serotyping with 7 unique O antigens and 13 H antigens detected and at least 14 different serotypes identified. The MAGs were also screened for virulence factor genes and as expected and similar to the ARGs, the *E*. *coli* MAGs had the greatest number of these genes (Table S8). Notably, among these *E*. *coli* MAGs was a predicted H49 serotype MAG (SUG3626) enriched in the d 4 florfenicol-treated piglets that was found to be carrying the *stx2* genes for Shiga toxin production. The *stx2* genes were from two variants, *stx2e* and *stx2f*. The heat-stable enterotoxin 1 (EAST-1) gene, *astA*, was found in two *E*. *coli* MAGs (SUG3618; SUG3629). The three *C*. *coli* MAGs carried the cytolethal distending toxin genes *cdtABC* while 10 *Campylobacter* sp945873855 MAGs as well as 2 *Helicobacter rappini* MAGs had only the *cdtB* gene. A *C*. *difficile* MAG (SUG3025) identified as sequence type 11 and enriched in the florfenicol-treated piglets on d 4 and 7, encoded two different *cdtA* and *cdtB* genes (*C*. *difficile* transferase) that share the same gene name but not function as the cytolethal distending toxin genes. In addition, genes for the major *C*. *difficile* virulence factors (*tcdA*/*toxA*) and toxin B (*tcdB*/*toxB*) (56) were also found in the *C*. *difficile* MAG here.

### Florfenicol-resistant isolates

To better understand the co-selection process, florfenicol-resistant bacteria (MIC > 32 µg/ml florfenicol) were isolated from fecal samples collected from florfenicol-treated piglets on d 4 and 11. These isolate genomes were then sequenced with both short- and long-read sequencing technologies. The florfenicol-resistant isolates recovered were classified as *E*. *coli* (n = 7), *Enterococcus avium* (n = 1), *E*. *faecalis* (n = 2), and *E*. *faecium* (n =1) (Table S9). The *E*. *coli* isolates were from three different serotypes: H49 (n = 3), O8:H25 (n = 3), and O4:H5 (n = 1). Using a 99.99% ANI threshold to define strains (57), there were 3 different strains among the 7 *E*. *coli* isolates, corresponding with the predicted serotypes (data not shown). Both *E*. *faecalis* isolates also appeared to belong to the same strain (> 99.99% ANI; data not shown). Genes conferring resistance to florfenicol were found in all 11 isolates with the enterococci carrying *fexB* and the *E*. *coli* isolates encoding *floR*. The *floR* gene was located on a plasmid together with *aph(3’’)-Ib*, *aph*(*6*)*-Id* (aminoglycosides), *bla*_TEM-1_ or *bla*_CMY-2_ (beta-lactams), *sul2* (sulfonamides), and *tet*(A) (tetracyclines) in the *E*. *coli* isolates (Fig. 8A, B). The *fexB* gene was co-located on a plasmid (23,095 to 27,535 bp) with *poxtA* in all four enterococci isolates (Fig. 8C,D).

**Figure 8.**
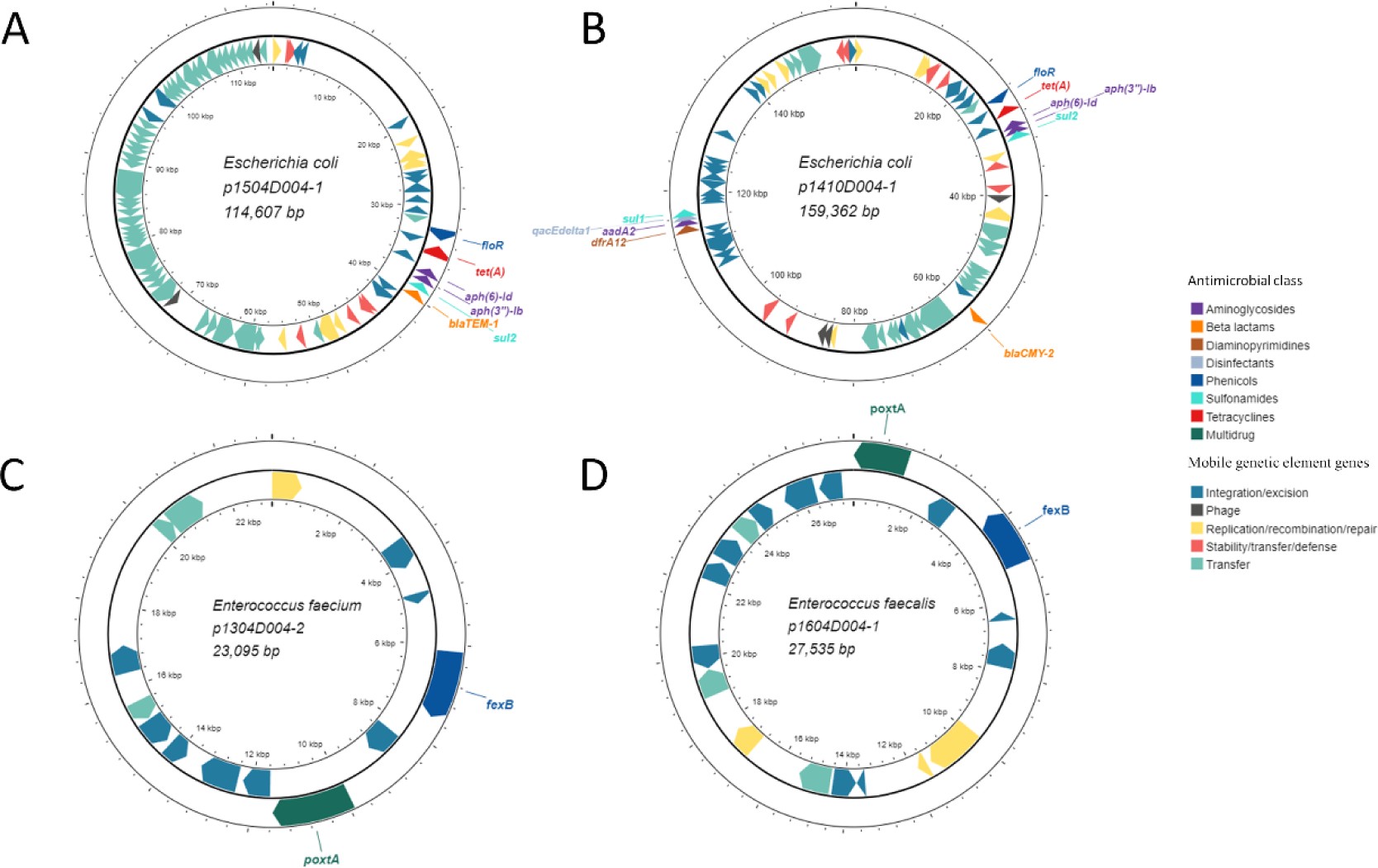
Locations of florfenicol resistance genes along with other antimicrobial resistance genes on plasmids in florfenicol-resistant *Escherichia coli* (A and B), *Enterococcus faecium* (C), and *Enterococcus faecalis* (D) isolates. The inner ring displays mobile genetic element genes colored by functional category.

This *fexB*-*poxtA* plasmid in both *E*. *avium* 1908D011 (p1908D011-2; 23,684 bp) and *E*. *faecium* 1304D004 (p1304D004-2; 23,095 bp) was 100% identical (100% coverage) to a plasmid (p1818-c; accession CP091209.1) in an *E*. *faecium* isolate from human feces in Switzerland (58) and was classified as a rep29 replicon (Fig. 8C; Table S9). *E*. *avium* 1908D011 also had a large plasmid (p1908D011-1; 129,571 bp) carrying *erm*(B) and *tet*(M) and with multiple replicons (repUS1, repUS43) that was 99.9% identical (47% coverage) to a plasmid in an *E*. *faecalis* strain (accession LR961926.1). Another plasmid (p1304D004-4; 30,712 bp) in *E*. *faecium* 1304D004 encoded *tet*(L), *tet*(M), *ant*(*6*)*-Ia*, *lnu*(B), and *lsa*(E). In the two *E*. *faecalis* isolates, the *fexB*-*poxtA* genes were on a multi-replicon (rep2, rep6) Inc18 plasmid (Fig. 8D; p1604D004-1, p1403D011-3) that was 100% identical (with 80% coverage) to a plasmid in an *E*. *faecium* strain recovered from cattle feces (59). To confirm that the *poxtA* gene was conferring the linezolid resistance phenotype, the MIC of linezolid for the four *Enterococcus* spp. isolates was determined. For both *E*. *faecalis* isolates, the MIC was 16 µg/ml and for *E*. *avium* 1908D011 and *E*. *faecium* the MIC of linezolid was 8 µg/ml. Based on CLSI breakpoints for linezolid and *Enterococcus*, these isolates were all resistant to linezolid. For the two ATCC strains without oxazolidinone resistance genes, *E*. *faecium* ATCC 35667 and *E*. *faecalis* ATCC 29212, the MIC of linezolid was 4 µg/ml.

In the six *E*. *coli* H49 and O8:H25 isolates, *floR*, *tet*(A), *aph*(*6*)*-Id*, *aph(3’’)-Ib*, *sul2*, and *bla*_TEM-1_ were co-located on a ≈ 114,600 bp IncI1-I(Alpha) plasmid (Fig. 8A). In *E*. *coli* 1410D004, *floR*, *tet*(A), *aph*(*6*)*-Id*, *aph(3’’)-Ib*, *sul2* were in the same order on a plasmid (p1410D004-1) as in the other six E. coli isolates; however, *bla*_TEM-1_ was replaced by *bla*_CMY-2_ which was also farther downstream of these other ARGs (Fig. 8B). In addition, p1410D004-1 was larger (159,362 bp) and contained another cluster of ARGs downstream: *dfrA12* (diaminopyrimidines), *aadA2* (aminoglycosides), *qacEdelta1* (disinfectants), and *sul1*. None of the *E*. *coli* isolates met the 99.99% ANI strain threshold with any of the *E*. *coli* MAGs, although the H49 *E*. *coli* isolates had 99.95% ANI with SUG3627 (data not shown). Both *E*. *faecalis* genomes had 99.5% ANI with *E*. *faecalis* SUG2336 but only 98.4% ANI with *E*. *faecalis* SUG2337. The genome of *E*. *faecium* 1304D004 exhibited 99.8% ANI with *E*. *faecium* SUG2339, while the ANI between *E*. *avium* 1908D011 and the *E*. *avium* MAG SUG2338 was 98.3% (data not shown).

## DISCUSSION

Despite 20 years of use in Canadian swine herds and longer in other countries, relatively little is known about how florfenicol affects the swine gut microbiome and resistome. Therefore, in this study we assessed the impact of florfenicol treatment in piglets on the development of their gut microbiome and resistome over the course of the swine production cycle. The gut microbiome of piglets treated with florfenicol at 1 day of age and again at 7 days of age differed significantly from that of the control piglets from days 4 through 14. These differences were at least partly driven by a consistent enrichment of *Clostridium* spp., *Escherichia* spp., and CALYPF01 sp945271245 in florfenicol-treated piglets and *B*. *rossii*, *Fusobacterium* spp., *P*. *hyovaginalis*, and *H. porci* in the control piglets. Although florfenicol had the greatest effect on the gut microbiome during the immediate periods following administration, there did appear to be some minor residual effects that persisted past the post-weaning phase, at least on day 84 of the study.

Given that florfenicol is a broad-spectrum antimicrobial active against a number of different gram-negative and gram-positive bacteria, it may be expected that multiple bacterial species would be affected in the immediate period following treatment. Furthermore, as previously noted, florfenicol’s usage in pigs is comparatively recent and is primarily administered through intramuscular injection, unlike chlortetracycline, lincomycin, or tylosin, which are typically added to feed for consecutive days or weeks and which have been used in pigs for many decades (60, 61). As a result, the pig gut microbiome, which is vertically transmitted from sows to piglets across generations, has had comparatively less exposure to florfenicol and therefore less time to adapt or develop resistance. The relatively quick recovery of the piglet gut microbiome to perturbation by florfenicol is likely due to the relatively short half-life of florfenicol (3) and the fact that the piglet gut microbiome has yet to be stably established prior to weaning.

Not unexpectedly, florfenicol treatment increased the relative abundance of the florfenicol resistance genes *fexA*, *fexB,* and *floR* as well as ARGs such as *cfr*, *cfr*(B), *clcD*, *optrA*, and *poxtA* that confer resistance to florfenicol and at least one other antimicrobial class (53). The enrichment of the *cfr*, *clcD, cfr*(B), *optrA*, and *poxtA* genes in treated piglets is particularly noteworthy as these genes also facilitate resistance to the oxazolidinones. Oxazolidinones are classified by the World Health Organization as critically important in human medicine (62) and include linezolid which is frequently used as an antibiotic of “last resort” against gram-positive multidrug-resistant bacteria such as methicillin-resistant *Staphylococcus aureus* and vancomycin-resistant enterococci (63). Although *poxtA* has not previously been reported in Canada, this is probably due to the fact that it was only first described in 2018 (54). The greater abundance of *fexB* and *floR* in the gut resistomes of florfenicol-treated piglets persisted until weaning, which was 14 days after the last treatment. Perhaps most significantly, florfenicol also increased the relative abundance of a large number of unrelated ARGs. Among these ARGs were those conferring resistance to aminoglycosides (*aadA8b*, *aph(3’’)-Ib*, *aph*(*6*)*-Id*), beta-lactams (*bla*_CMY-59_), diaminopyrimidines (*dfrA12*), sulfonamides (*sul2*), and tetracyclines [*tet*(A)]. Drugs within these antimicrobial classes are all used in swine in Canada via feed, injection, or water at varying frequencies (60, 61) and this likely explains why these ARGs initially emerged in the swine gut.

The MAGs recovered from the metagenomes were screened for ARGs to provide taxonomic context. In terms of florfenicol resistance genes, a *V*. *lutrae* MAG was identified carrying *fexA* together with *ant(4’)-Ib* and *optrA* while a *S*. *borealis* MAG also carried *fexA*. However, neither the MAGs nor the bacterial species were differentially abundant between the two groups of pigs. Notably, multidrug-resistant *S*. *borealis* isolates with *fexA* and several other ARGs have recently been recovered from the nasal cavity of pigs (64). Similarly, *fexA* and *optrA* have been identified on the chromosome of *V*. *lutrae* isolated from the lung of a pig (65) and human feces (66). The *poxtA* gene, which is reported to provide resistance to tetracyclines in addition to phenicols and oxazolidinones (67), was detected in MAGs that were classified as *Anaerotignum* sp001304995, *Eisenbergiella porci*, *Enterocloster porci*, *E. clostridioformis*, or *H. hathewayi* and that were relatively more abundant in the metagenomes of florfenicol-treated piglets at d 4 or 7. This is in agreement with reports of *poxtA* in *H*. *hathewayi* (68) and *E*. *clostridioformis* (54) isolate genomes.

Overall, ARGs conferring resistance to tetracyclines and MLS_B_ remained largely unaffected by florfenicol treatment. However, there were certain ARGs within these classes that had a higher relative abundance in the control piglets at specific time points. Some of these ARGs such as *lnu*(C), *tet*(B), *te*t(T), *tet*(Q), *tet*(W/32/O) were also binned into MAGs that were enriched in the control piglets. For example, three *P*. *hyovaginalis* MAGs that were relatively more abundant in the metagenomes of the control pigs carried *tet*(W) and/or *tet*(W/32/O). Additionally, *tet*(Q) was binned into 27 *Collinsella* sp002391315 MAGs that were all enriched in the control piglet microbiomes on day 4. This is consistent with reports of *tet* genes in certain *Collinsella* spp. strains (69). Similarly, two *Basfia porcinus* MAGs carrying *tet*(B) were also relatively more abundant in the untreated pigs on days 4, 11, and 14. The type strain for this species (NCBI: *Actinobacillus porcinus* NM319), originally isolated from the pig respiratory tract, was reported to carry the *tet*(B) gene (70). It is therefore clear from this and earlier studies, that ARGs conferring resistance to the MLS_B_ and tetracycline classes persist even in the absence of exposure to these antimicrobials (8, 15, 71, 72).

Although MAGs can provide genomic and taxonomic context for certain chromosomally-encoded ARGs, many ARGs are not binned into MAGs, especially those on MGEs such as plasmids (73). Therefore, we isolated and sequenced the genomes of bacteria displaying phenotypic resistance to florfenicol (MIC > 32 µg/ml) that were recovered from the feces of treated piglets. The majority of these isolates were identified as *E*. *coli* (n = 7) from 3 different serotypes. All seven *E*. *coli* isolates carried *floR* and most importantly, this gene was co-located on a plasmid with *aph(3’’)-Ib*, *aph*(*6*)*-Id*, *bla*_TEM-1_/*bla*_CMY-2_, *sul2*, and *tet*(A). This likely explains the enrichment of these ARGs in the resistomes of piglets administered florfenicol particularly since *E*. *coli* and other *Escherichia* spp. were also relatively more abundant in these piglets. Plasmids carrying these same ARGs have been identified in *E*. *coli* isolated from corvids (74), humans (75), pigs (76), dairy cattle manure (77), poultry (78), and groundwater (79). These ARGs have also been found on plasmids in other *Enterobacteriaceae* including *Salmonella enterica* subsp. *enterica* recovered from beef cattle (80) and their environment (81), as well as horses (80), humans (82), pigs (80, 83), and poultry (80), plus *Proteus mirabilis* from humans (84, 85). Thus, these ARGs appear to be widely disseminated together across multiple hosts and environments. Furthermore, it is probable that an antimicrobial from any of these classes may co-select for all of these ARGs.

The four enterococci isolates recovered from the feces of the florfenicol-treated piglets carried *poxtA* (and *fexB*) and were confirmed to be phenotypically resistant to linezolid. As with *floR* and other ARGs in the *E*. *coli* isolates, the *fexB* and *poxtA* genes were co-located on a plasmid in the *E*. *avium*, *E*. *faecalis*, and *E*. *faecium* isolates. The *fexB*-*poxtA* plasmid was nearly identical to ones found in *E. faecium* isolates from cattle (59), humans (58, 86), and pigs (87, 88) in Europe and China indicating wide geographic and host distribution. Although nominally the *poxtA* confers resistance to tetracyclines it is unclear whether this is true (67, 89). Additionally, oxazolidinones have never been approved for use in food-producing animals and therefore the presence of two co-located ARGs conferring resistance to phenicols is notable. The *fexB* and *poxtA* genes mediate phenicol resistance through distinct mechanisms, suggesting that the simultaneous presence of both genes results in higher levels of resistance to phenicols compared to the presence of only one gene.

Another objective of this study was to assess the potential transfer of bacteria and ARGs from the sows to their piglets. The piglet gut microbiome is initially seeded with microbes from the vaginal tract during birth and from the colostrum they consume in the immediate postnatal period. Piglets are coprophagic and as such, feces from the sow are also a significant source of bacteria for the piglet gut. In the present study, using source tracking it was predicted that colostrum was the largest contributor to the piglet gut microbiome in the immediate post-farrowing period. This may be expected as piglets consume colostrum soon after birth and this represents one of their first exposures to microorganisms. Many of the most abundant microbial species identified in the colostrum such as *Methanobrevibacter* sp900769095, *Sodaliphilus* sp004557565, *Prevotella* sp900546535 [recently proposed name: *Segatella brasiliensis* (90)], and *Prevotella* sp000434975 are associated with the pig (91–93) or human gut (90). Therefore, although the teat was cleaned prior to sample collection it is likely that the colostrum samples contained skin and fecal bacteria. Nonetheless, these are bacteria that the piglets would naturally be exposed to while suckling on the teat. The largest contributor to the piglet resistome was predicted to be the sow’s feces, with post-weaned pigs showing considerable overlap in their resistome with that of the sows. As mentioned, this was particularly evident for genes conferring resistance to the tetracycline and MLS_B_ classes.

In summary, treatment with florfenicol in early life increased the abundance of ARGs conferring resistance to not only the phenicols but to several other antimicrobial classes as well. Furthermore, *E*. *coli* isolated from the feces of treated piglets contained a large plasmid with *floR* and other co-located ARGs providing resistance to at least four other antimicrobial classes. In addition, enterococci isolates carried *poxtA*, a multidrug resistance gene, on a plasmid together with *fexB*. Thus, this study shows the potential risk that florfenicol use may pose in terms of co-selection and transfer of multiple ARGs. Although the effect on the piglet gut microbiome and resistome was largely lost post-weaning, it would be interesting to know if florfenicol would have these same effects if administered post-weaning.

## Acknowledgements

The authors would like to thank Cara Service and the animal care staff for their excellent care of the pigs and for their help with sampling as well. We are also grateful to Chloe Brennan, Rylan Lacey, Tingting Liu, Sean Norris, and Quinn Tomalty for their help with laboratory work. The authors also appreciate helpful feedback from Cassidy Klima. Funding for this project was provided by Agriculture and Agri-Food Canada’s A-Base program.

## Author contributions

D.B.H. designed the research. A.K., D.B.H., and K.E.G. carried out bioinformatics and statistical analyses. K.E.G. conducted the laboratory experiments. D.B.H. wrote the manuscript. A.K. and K.E.G. reviewed and edited the manuscript. All authors read and approved the final manuscript.

## Data availability

All metagenomic sequences, metagenome-assembled genomes, and isolate genomes are publicly available in the National Center for Biotechnology Information’s (NCBI) sequence read archive and genome databases under BioProject PRJNA779404.

## References

1. Schwarz S, Kehrenberg C, Doublet B, Cloeckaert A. 2004. Molecular basis of bacterial resistance to chloramphenicol and florfenicol. FEMS Microbiol Rev 28:519–42. 10.1016/j.femsre.2004.04.001.

2. Somogyi Z, Mag P, Kovacs D, Kerek A, Szabo P, Makrai L, Jerzsele A. 2022. Synovial and systemic pharmacokinetics of florfenicol and PK/PD integration against *Streptococcus suis* in pigs. Pharmaceutics 14:109. 10.3390/pharmaceutics14010109.

3. Dorey L, Pelligand L, Cheng Z, Lees P. 2017. Pharmacokinetic/pharmacodynamic integration and modelling of florfenicol for the pig pneumonia pathogens *Actinobacillus pleuropneumoniae* and *Pasteurella multocida*. PLoS One 12:e0177568. 10.1371/journal.pone.0177568.

4. De Smet J, Boyen F, Croubels S, Rasschaert G, Haesebrouck F, De Backer P, Devreese M. 2018. Similar gastro-intestinal exposure to florfenicol after oral or intramuscular administration in pigs, leading to resistance selection in commensal *Escherichia coli*. Front Pharmacol 9:1265. 10.3389/fphar.2018.01265.

5. Liu H, Wang Y, Wu C, Schwarz S, Shen Z, Jeon B, Ding S, Zhang Q, Shen J. 2012. A novel phenicol exporter gene, *fexB*, found in enterococci of animal origin. J Antimicrob Chemother 67:322–5. 10.1093/jac/dkr481.

6. Hao W, Shan X, Li D, Schwarz S, Zhang SM, Li XS, Du XD. 2019. Analysis of a *poxtA*- and *optrA*-co-carrying conjugative multiresistance plasmid from *Enterococcus faecalis*. J Antimicrob Chemother 74:1771–1775. 10.1093/jac/dkz109.

7. Partridge SR, Kwong SM, Firth N, Jensen SO. 2018. Mobile genetic elements associated with antimicrobial resistance. Clin Microbiol Rev 31:10.1128/cmr.00088-17. 10.1128/CMR.00088-17.

8. Holman DB, Gzyl KE, Mou KT, Allen HK. 2021. Weaning age and its effect on the development of the swine gut microbiome and resistome. mSystems 6:e0068221. 10.1128/mSystems.00682-21.

9. Holman DB, Gzyl KE, Scott H, Cara S, Prieto N, Lopez-Campos O. 2024. Associations between the rumen microbiota and carcass merit and meat quality in beef cattle. Appl Microbiol Biotechnol 108:287. 10.1007/s00253-024-13126-1.

10. Martin M. 2011. Cutadapt removes adapter sequences from high-throughput sequencing reads. EMBnet journal 17:10–12. 10.14806/ej.17.1.200.

11. Callahan BJ, McMurdie PJ, Rosen MJ, Han AW, Johnson AJ, Holmes SP. 2016. DADA2: High-resolution sample inference from Illumina amplicon data. Nat Methods 13:581–3. 10.1038/nmeth.3869.

12. Quast C, Pruesse E, Yilmaz P, Gerken J, Schweer T, Yarza P, Peplies J, Glockner FO. 2013. The SILVA ribosomal RNA gene database project: improved data processing and web-based tools. Nucleic Acids Res 41:D590–6. 10.1093/nar/gks1219.

13. McMurdie PJ, Holmes S. 2013. phyloseq: an R package for reproducible interactive analysis and graphics of microbiome census data. PLoS One 8:e61217. 10.1371/journal.pone.0061217.

14. Oksanen J, Simpson GL, Blanchet FG, Kindt R, Legendre P, Minchin PR, O’Hara R, Solymos P, Stevens M, Szoecs E. 2023. Vegan: community ecology package, 2.6-4. Vienna (Austria): R Foundation for Statistical Computing.

15. Holman DB, Gzyl KE, Kommadath A. 2023. The gut microbiome and resistome of conventionally vs. pasture-raised pigs. Microb Genom 9:2023.03. 02.530897. 10.1099/mgen.0.001061.

16. Chen S, Zhou Y, Chen Y, Gu J. 2018. fastp: an ultra-fast all-in-one FASTQ preprocessor. Bioinformatics 34:i884–i890. 10.1093/bioinformatics/bty560.

17. Langmead B, Salzberg SL. 2012. Fast gapped-read alignment with Bowtie 2. Nat Methods 9:357–9. 10.1038/nmeth.1923.

18. Li H, Handsaker B, Wysoker A, Fennell T, Ruan J, Homer N, Marth G, Abecasis G, Durbin R, Genome Project Data Processing S. 2009. The sequence alignment/map format and SAMtools. Bioinformatics 25:2078–9. 10.1093/bioinformatics/btp352.

19. Quinlan AR. 2014. BEDTools: The Swiss-Army Tool for Genome Feature Analysis. Curr Protoc Bioinformatics 47:11 12 1–34. 10.1002/0471250953.bi1112s47.

20. Wood DE, Lu J, Langmead B. 2019. Improved metagenomic analysis with Kraken 2. Genome Biol 20:257. 10.1186/s13059-019-1891-0.

21. Lu J, Breitwieser FP, Thielen P, Salzberg SL. 2017. Bracken: estimating species abundance in metagenomics data. PeerJ Computer Science 3:e104. 10.7717/peerj-cs.104.

22. Parks DH, Chuvochina M, Rinke C, Mussig AJ, Chaumeil PA, Hugenholtz P. 2022. GTDB: an ongoing census of bacterial and archaeal diversity through a phylogenetically consistent, rank normalized and complete genome-based taxonomy. Nucleic Acids Res 50:D785–D794. 10.1093/nar/gkab776.

23. Alcock BP, Huynh W, Chalil R, Smith KW, Raphenya AR, Wlodarski MA, Edalatmand A, Petkau A, Syed SA, Tsang KK, Baker SJC, Dave M, McCarthy MC, Mukiri KM, Nasir JA, Golbon B, Imtiaz H, Jiang X, Kaur K, Kwong M, Liang ZC, Niu KC, Shan P, Yang JYJ, Gray KL, Hoad GR, Jia B, Bhando T, Carfrae LA, Farha MA, French S, Gordzevich R, Rachwalski K, Tu MM, Bordeleau E, Dooley D, Griffiths E, Zubyk HL, Brown ED, Maguire F, Beiko RG, Hsiao WWL, Brinkman FSL, Van Domselaar G, McArthur AG. 2023. CARD 2023: expanded curation, support for machine learning, and resistome prediction at the Comprehensive Antibiotic Resistance Database. Nucleic Acids Res 51:D690–D699. 10.1093/nar/gkac920.

24. Clausen P, Aarestrup FM, Lund O. 2018. Rapid and precise alignment of raw reads against redundant databases with KMA. BMC Bioinformatics 19:307. 10.1186/s12859-018-2336-6.

25. Li D, Liu CM, Luo R, Sadakane K, Lam TW. 2015. MEGAHIT: an ultra-fast single-node solution for large and complex metagenomics assembly via succinct de Bruijn graph. Bioinformatics 31:1674–6. 10.1093/bioinformatics/btv033.

26. Kang DD, Li F, Kirton E, Thomas A, Egan R, An H, Wang Z. 2019. MetaBAT 2: an adaptive binning algorithm for robust and efficient genome reconstruction from metagenome assemblies. PeerJ 7:e7359. 10.7717/peerj.7359.

27. Parks DH, Imelfort M, Skennerton CT, Hugenholtz P, Tyson GW. 2015. CheckM: assessing the quality of microbial genomes recovered from isolates, single cells, and metagenomes. Genome Res 25:1043–55. 10.1101/gr.186072.114.

28. Chaumeil PA, Mussig AJ, Hugenholtz P, Parks DH. 2022. GTDB-Tk v2: memory friendly classification with the genome taxonomy database. Bioinformatics 38:5315–5316. 10.1093/bioinformatics/btac672.

29. Buchfink B, Reuter K, Drost HG. 2021. Sensitive protein alignments at tree-of-life scale using DIAMOND. Nat Methods 18:366–368. 10.1038/s41592-021-01101-x.

30. Liu B, Zheng D, Zhou S, Chen L, Yang J. 2022. VFDB 2022: a general classification scheme for bacterial virulence factors. Nucleic Acids Res 50:D912–D917. 10.1093/nar/gkab1107.

31. Joensen KG, Tetzschner AM, Iguchi A, Aarestrup FM, Scheutz F. 2015. Rapid and easy in silico serotyping of *Escherichia coli* isolates by use of whole-genome sequencing data. J Clin Microbiol 53:2410–26. 10.1128/JCM.00008-15.

32. Jolley KA, Bray JE, Maiden MCJ. 2018. Open-access bacterial population genomics: BIGSdb software, the PubMLST.org website and their applications. Wellcome Open Res 3:124. 10.12688/wellcomeopenres.14826.1.

33. Asnicar F, Thomas AM, Beghini F, Mengoni C, Manara S, Manghi P, Zhu Q, Bolzan M, Cumbo F, May U, Sanders JG, Zolfo M, Kopylova E, Pasolli E, Knight R, Mirarab S, Huttenhower C, Segata N. 2020. Precise phylogenetic analysis of microbial isolates and genomes from metagenomes using PhyloPhlAn 3.0. Nat Commun 11:2500. 10.1038/s41467-020-16366-7.

34. Li J, Shao B, Shen J, Wang S, Wu Y. 2013. Occurrence of chloramphenicol-resistance genes as environmental pollutants from swine feedlots. Environ Sci Technol 47:2892–7. 10.1021/es304616c.

35. Wick RR, Judd LM, Holt KE. 2023. Assembling the perfect bacterial genome using Oxford Nanopore and Illumina sequencing. PLoS Comput Biol 19:e1010905. 10.1371/journal.pcbi.1010905.

36. Wick RR, Judd LM, Cerdeira LT, Hawkey J, Meric G, Vezina B, Wyres KL, Holt KE. 2021. Trycycler: consensus long-read assemblies for bacterial genomes. Genome Biol 22:266. 10.1186/s13059-021-02483-z.

37. Kolmogorov M, Yuan J, Lin Y, Pevzner PA. 2019. Assembly of long, error-prone reads using repeat graphs. Nat Biotechnol 37:540–546. 10.1038/s41587-019-0072-8.

38. Li H. 2016. Minimap and miniasm: fast mapping and de novo assembly for noisy long sequences. Bioinformatics 32:2103–10. 10.1093/bioinformatics/btw152.

39. Li H. 2018. Minimap2: pairwise alignment for nucleotide sequences. Bioinformatics 34:3094–3100. 10.1093/bioinformatics/bty191.

40. Wick RR, Holt KE. 2019. Benchmarking of long-read assemblers for prokaryote whole genome sequencing. F1000Res 8:2138. 10.12688/f1000research.21782.4.

41. Vaser R, Sikic M. 2021. Time- and memory-efficient genome assembly with Raven. Nat Comput Sci 1:332–336. 10.1038/s43588-021-00073-4.

42. Vaser R, Sovic I, Nagarajan N, Sikic M. 2017. Fast and accurate de novo genome assembly from long uncorrected reads. Genome Res 27:737–746. 10.1101/gr.214270.116.

43. Koren S, Walenz BP, Berlin K, Miller JR, Bergman NH, Phillippy AM. 2017. Canu: scalable and accurate long-read assembly via adaptive k-mer weighting and repeat separation. Genome Res 27:722–736. 10.1101/gr.215087.116.

44. Wick RR, Holt KE. 2022. Polypolish: Short-read polishing of long-read bacterial genome assemblies. PLoS Comput Biol 18:e1009802. 10.1371/journal.pcbi.1009802.

45. Li H, Durbin R. 2009. Fast and accurate short read alignment with Burrows-Wheeler transform. Bioinformatics 25:1754–60. 10.1093/bioinformatics/btp324.

46. Chklovski A, Parks DH, Woodcroft BJ, Tyson GW. 2023. CheckM2: a rapid, scalable and accurate tool for assessing microbial genome quality using machine learning. Nat Methods 20:1203–1212. 10.1038/s41592-023-01940-w.

47. Carattoli A, Zankari E, Garcia-Fernandez A, Voldby Larsen M, Lund O, Villa L, Moller Aarestrup F, Hasman H. 2014. In silico detection and typing of plasmids using PlasmidFinder and plasmid multilocus sequence typing. Antimicrob Agents Chemother 58:3895–903. 10.1128/AAC.02412-14.

48. Jain C, Rodriguez RL, Phillippy AM, Konstantinidis KT, Aluru S. 2018. High throughput ANI analysis of 90K prokaryotic genomes reveals clear species boundaries. Nat Commun 9:5114. 10.1038/s41467-018-07641-9.

49. Grant JR, Enns E, Marinier E, Mandal A, Herman EK, Chen CY, Graham M, Van Domselaar G, Stothard P. 2023. Proksee: in-depth characterization and visualization of bacterial genomes. Nucleic Acids Res 51:W484–W492. 10.1093/nar/gkad326.

50. Mallick H, Rahnavard A, McIver LJ, Ma S, Zhang Y, Nguyen LH, Tickle TL, Weingart G, Ren B, Schwager EH, Chatterjee S, Thompson KN, Wilkinson JE, Subramanian A, Lu Y, Waldron L, Paulson JN, Franzosa EA, Bravo HC, Huttenhower C. 2021. Multivariable association discovery in population-scale meta-omics studies. PLoS Comput Biol 17:e1009442. 10.1371/journal.pcbi.1009442.

51. Bates D, Maechler M, Bolker B, Walker S. 2014. lme4: Linear mixed-effects models using Eigen and S4. R package version 1.1–7.

52. Knights D, Kuczynski J, Charlson ES, Zaneveld J, Mozer MC, Collman RG, Bushman FD, Knight R, Kelley ST. 2011. Bayesian community-wide culture-independent microbial source tracking. Nat Methods 8:761–3. 10.1038/nmeth.1650.

53. Long KS, Poehlsgaard J, Kehrenberg C, Schwarz S, Vester B. 2006. The Cfr rRNA methyltransferase confers resistance to phenicols, lincosamides, oxazolidinones, pleuromutilins, and streptogramin A antibiotics. Antimicrob Agents Chemother 50:2500–5. 10.1128/AAC.00131-06.

54. Antonelli A, D’Andrea MM, Brenciani A, Galeotti CL, Morroni G, Pollini S, Varaldo PE, Rossolini GM. 2018. Characterization of *poxtA*, a novel phenicol-oxazolidinone-tetracycline resistance gene from an MRSA of clinical origin. J Antimicrob Chemother 73:1763–1769. 10.1093/jac/dky088.

55. Peltier J, Courtin P, El Meouche I, Catel-Ferreira M, Chapot-Chartier MP, Lemee L, Pons JL. 2013. Genomic and expression analysis of the *vanG*-like gene cluster of *Clostridium difficile*. Microbiology (Reading) 159:1510–1520. 10.1099/mic.0.065060-0.

56. Kordus SL, Thomas AK, Lacy DB. 2022. *Clostridioides difficile* toxins: mechanisms of action and antitoxin therapeutics. Nat Rev Microbiol 20:285–298. 10.1038/s41579-021-00660-2.

57. Rodriguez RL, Conrad RE, Viver T, Feistel DJ, Lindner BG, Venter SN, Orellana LH, Amann R, Rossello-Mora R, Konstantinidis KT. 2024. An ANI gap within bacterial species that advances the definitions of intra-species units. mBio 15:e0269623. 10.1128/mbio.02696-23.

58. Nuesch-Inderbinen M, Biggel M, Zurfluh K, Treier A, Stephan R. 2022. Faecal carriage of enterococci harbouring oxazolidinone resistance genes among healthy humans in the community in Switzerland. J Antimicrob Chemother 77:2779–2783. 10.1093/jac/dkac260.

59. Nuesch-Inderbinen M, Biggel M, Haussmann A, Treier A, Heyvaert L, Cernela N, Stephan R. 2023. Oxazolidinone resistance genes in florfenicol-resistant enterococci from beef cattle and veal calves at slaughter. Front Microbiol 14:1150070. 10.3389/fmicb.2023.1150070.

60. Bosman AL, Deckert AE, Carson CA, Poljak Z, Reid-Smith RJ, McEwen SA. 2022. Antimicrobial use in lactating sows, piglets, nursery, and grower-finisher pigs on swine farms in Ontario, Canada during 2017 and 2018. Porcine Health Manag 8:17. 10.1186/s40813-022-00259-w.

61. Government of Canada. 2022. CIPARS: Canadian integrated program for antimicrobial resistance surveillance - pigs. Public Health Agency of Canada. Guelph. ON. Accessed: May 31, 2024. https://publications.gc.ca/collections/collection_2022/aspc-phac/HP2-4-2019-eng-5.pdf.

62. World Health Organization. 2019. Critically important antimicrobials for human medicine. Switzerland. Accessed: May 31, 2024. https://iris.who.int/bitstream/handle/10665/312266/9789241515528-eng.pdf

63. Zahedi Bialvaei A, Rahbar M, Yousefi M, Asgharzadeh M, Samadi Kafil H. 2017. Linezolid: a promising option in the treatment of Gram-positives. J Antimicrob Chemother 72:354–364. 10.1093/jac/dkw450.

64. Abdullahi IN, Lozano C, Simon C, Zarazaga M, Torres C. 2023. Within-host diversity of coagulase-negative staphylococci resistome from healthy pigs and pig farmers, with the detection of *cfr*-carrying strains and MDR-*S*. *borealis*. Antibiotics (Basel) 12:1505. 10.3390/antibiotics12101505.

65. Zhu Y, Yang W, Schwarz S, Xu Q, Yang Q, Wang L, Liu S, Zhang W. 2022. Characterization of the novel *optrA*-carrying pseudo-compound transposon Tn7363 and an Inc18 plasmid carrying *cfr*(D) in *Vagococcus lutrae*. J Antimicrob Chemother 77:921–925. 10.1093/jac/dkab478.

66. Shen W, Huang Y, Cai J. 2022. An optimized screening approach for the oxazolidinone resistance gene *optrA* yielded a higher fecal carriage rate among healthy individuals in Hangzhou, China. Microbiol Spectr 10:e0297422. 10.1128/spectrum.02974-22.

67. Crowe-McAuliffe C, Murina V, Turnbull KJ, Huch S, Kasari M, Takada H, Nersisyan L, Sundsfjord A, Hegstad K, Atkinson GC, Pelechano V, Wilson DN, Hauryliuk V. 2022. Structural basis for PoxtA-mediated resistance to phenicol and oxazolidinone antibiotics. Nat Commun 13:1860. 10.1038/s41467-022-29274-9.

68. Hernández-Juárez LE, Camorlinga M, Méndez-Tenorio A, Calderón JF, Huang BC, Bandoy DDR, Weimer BC, Torres J. 2021. Analyses of publicly available *Hungatella hathewayi* genomes revealed genetic distances indicating they belong to more than one species. Virulence 12:1950–1964.

69. Bag S, Ghosh TS, Das B. 2017. Complete Genome Sequence of Collinsella aerofaciens Isolated from the Gut of a Healthy Indian Subject. Genome Announc 5:10.1128/genomea.01361-17. 10.1128/genomeA.01361-17.

70. Bosse JT, Li Y, Fernandez Crespo R, Angen O, Holden MTG, Weinert LA, Maskell DJ, Tucker AW, Wren BW, Rycroft AN, Langford PR, consortium BRT. 2020. Draft genome sequences of the type strains of *Actinobacillus indolicus* (46K2C) and *Actinobacillus porcinus* (NM319), two NAD-dependent bacterial species found in the respiratory tract of pigs. Microbiol Resour Announc 9:10.1128/mra.00716-19. 10.1128/MRA.00716-19.

71. Pollock J, Muwonge A, Hutchings MR, Mainda G, Bronsvoort BM, Gally DL, Corbishley A. 2020. Resistance to change: AMR gene dynamics on a commercial pig farm with high antimicrobial usage. Sci Rep 10:1708. 10.1038/s41598-020-58659-3.

72. Brinck JE, Lassen SB, Forouzandeh A, Pan T, Wang YZ, Monteiro A, Blavi L, Sola-Oriol D, Stein HH, Su JQ, Brandt KK. 2023. Impacts of dietary copper on the swine gut microbiome and antibiotic resistome. Sci Total Environ 857:159609. 10.1016/j.scitotenv.2022.159609.

73. Maguire F, Jia B, Gray KL, Lau WYV, Beiko RG, Brinkman FSL. 2020. Metagenome-assembled genome binning methods with short reads disproportionately fail for plasmids and genomic Islands. Microb Genom 6. 10.1099/mgen.0.000436.

74. Jamborova I, Janecko N, Halova D, Sedmik J, Mezerova K, Papousek I, Kutilova I, Dolejska M, Cizek A, Literak I. 2018. Molecular characterization of plasmid-mediated AmpC beta-lactamase- and extended-spectrum beta-lactamase-producing *Escherichia coli* and *Klebsiella pneumoniae* among corvids (*Corvus brachyrhynchos* and *Corvus corax*) roosting in Canada. FEMS Microbiol Ecol 94:fiy166. 10.1093/femsec/fiy166.

75. Day M, Doumith M, Jenkins C, Dallman TJ, Hopkins KL, Elson R, Godbole G, Woodford N. 2017. Antimicrobial resistance in *Shiga* toxin-producing *Escherichia coli* serogroups O157 and O26 isolated from human cases of diarrhoeal disease in England, 2015. J Antimicrob Chemother 72:145–152. 10.1093/jac/dkw371.

76. Hayer SS, Lim S, Hong S, Elnekave E, Johnson T, Rovira A, Vannucci F, Clayton JB, Perez A, Alvarez J. 2020. Genetic determinants of resistance to extended-spectrum cephalosporin and fluoroquinolone in *Escherichia coli* isolated from diseased pigs in the United States. mSphere 5. 10.1128/mSphere.00990-20.

77. Anderson REV, Chalmers G, Murray R, Mataseje L, Pearl DL, Mulvey M, Topp E, Boerlin P. 2023. Characterization of *Escherichia coli* and other Enterobacterales resistant to extended-spectrum cephalosporins isolated from dairy manure in Ontario, Canada. Appl Environ Microbiol 89:e0186922. 10.1128/aem.01869-22.

78. Rehman MA, Rempel H, Carrillo CD, Ziebell K, Allen K, Manges AR, Topp E, Diarra MS. 2022. Virulence genotype and phenotype of multiple antimicrobial-resistant *Escherichia coli* Isolates from broilers assessed from a “One-Health” perspective. J Food Prot 85:336–354. 10.4315/JFP-21-273.

79. Andrade L, M PR, L PB, Hynds P, Weatherill J, O’Dwyer J. 2023. Assessing antimicrobial and metal resistance genes in *Escherichia coli* from domestic groundwater supplies in rural Ireland. Environ Pollut 333:121970. 10.1016/j.envpol.2023.121970.

80. McMillan EA, Gupta SK, Williams LE, Jove T, Hiott LM, Woodley TA, Barrett JB, Jackson CR, Wasilenko JL, Simmons M, Tillman GE, McClelland M, Frye JG. 2019. Antimicrobial resistance genes, cassettes, and plasmids present in *Salmonella enterica* associated with United States food animals. Front Microbiol 10:832. 10.3389/fmicb.2019.00832.

81. Lewis GL, Fenton RJ, Moriyama EN, Loy JD, Moxley RA. 2023. Association of ISVsa3 with multidrug resistance in *Salmonella enterica* isolates from cattle (*Bos taurus*). Microorganisms 11:631. 10.3390/microorganisms11030631.

82. Liu M, Zhu K, Li X, Han Y, Yang C, Liu H, Du X, Xu X, Yang H, Song H, Qiu S, Xiang Y. 2023. Genetic characterization of a *Salmonella enterica* serovar Typhimurium isolated from an infant with concurrent resistance to ceftriaxone, ciprofloxacin and azithromycin. J Glob Antimicrob Resist 35:252–256. 10.1016/j.jgar.2023.09.016.

83. Ge B, Mukherjee S, Li C, Harrison LB, Hsu CH, Tran TT, Whichard JM, Dessai U, Singh R, Gilbert JM, Strain EA, McDermott PF, Zhao S. 2024. Genomic analysis of azithromycin-resistant *Salmonella* from food animals at slaughter and processing, and retail meats, 2011-2021, United States. Microbiol Spectr 12:e0348523. 10.1128/spectrum.03485-23.

84. Osei Sekyere J, Maningi NE, Modipane L, Mbelle NM. 2020. Emergence of *mcr-9.1* in extended-spectrum-beta-lactamase-producing clinical *Enterobacteriaceae* in Pretoria, South Africa: global evolutionary phylogenomics, resistome, and mobilome. mSystems 5. 10.1128/mSystems.00148-20.

85. Johnning A, Karami N, Tang Hallback E, Muller V, Nyberg L, Buongermino Pereira M, Stewart C, Ambjornsson T, Westerlund F, Adlerberth I, Kristiansson E. 2018. The resistomes of six carbapenem-resistant pathogens - a critical genotype-phenotype analysis. Microb Genom 4:e000233. 10.1099/mgen.0.000233.

86. Dejoies L, Sassi M, Schutz S, Moreaux J, Zouari A, Potrel S, Collet A, Lecourt M, Auger G, Cattoir V. 2021. Genetic features of the *poxtA* linezolid resistance gene in human enterococci from France. J Antimicrob Chemother 76:1978–1985. 10.1093/jac/dkab116.

87. Shan X, Li XS, Wang N, Schwarz S, Zhang SM, Li D, Du XD. 2020. Studies on the role of IS1216E in the formation and dissemination of *poxtA*-carrying plasmids in an *Enterococcus faecium* clade A1 isolate. J Antimicrob Chemother 75:3126–3130. 10.1093/jac/dkaa325.

88. Nuesch-Inderbinen M, Haussmann A, Treier A, Zurfluh K, Biggel M, Stephan R. 2022. Fattening pigs are a reservoir of florfenicol-resistant enterococci harboring oxazolidinone resistance genes. J Food Prot 85:740–746. 10.4315/JFP-21-431.

89. Freitas AR, Tedim AP, Duarte B, Elghaieb H, Abbassi MS, Hassen A, Read A, Alves V, Novais C, Peixe L. 2020. Linezolid-resistant (Tn6246::*fexB*-*poxtA*) *Enterococcus faecium* strains colonizing humans and bovines on different continents: similarity without epidemiological link. J Antimicrob Chemother 75:2416–2423. 10.1093/jac/dkaa227.

90. Blanco-Miguez A, Galvez EJC, Pasolli E, De Filippis F, Amend L, Huang KD, Manghi P, Lesker TR, Riedel T, Cova L, Puncochar M, Thomas AM, Valles-Colomer M, Schober I, Hitch TCA, Clavel T, Berry SE, Davies R, Wolf J, Spector TD, Overmann J, Tett A, Ercolini D, Segata N, Strowig T. 2023. Extension of the *Segatella copri* complex to 13 species with distinct large extrachromosomal elements and associations with host conditions. Cell Host Microbe 31:1804–1819 e9. 10.1016/j.chom.2023.09.013.

91. Holman DB, Kommadath A, Tingley JP, Abbott DW. 2022. Novel insights into the pig gut microbiome using metagenome-assembled genomes. Microbiol Spectr 10:e0238022. 10.1128/spectrum.02380-22.

92. Gaio D, DeMaere MZ, Anantanawat K, Chapman TA, Djordjevic SP, Darling AE. 2021. Post-weaning shifts in microbiome composition and metabolism revealed by over 25 000 pig gut metagenome-assembled genomes. Microb Genom 7. 10.1099/mgen.0.000501.

93. Crossfield M, Gilroy R, Ravi A, Baker D, La Ragione RM, Pallen MJ. 2022. Archaeal and bacterial metagenome-assembled genome sequences derived from pig feces. Microbiol Resour Announc 11:e0114221. 10.1128/mra.01142-21.

